# Variations in the latitudinal diversity gradients of the ocean microbiome

**DOI:** 10.1101/2025.10.13.682024

**Authors:** Dominic Eriksson, Jonas Schiller, Alexandre Schickele, Taylor Priest, Anna Mankowski, Enzo Faucher, Lucas J. Ustick, Michael Kuhn, Samuel Miravet-Verde, Hans-Joachim Ruscheweyh, Corentin Clerc, Nicolas Gruber, Shinichi Sunagawa, Peer Bork, Meike Vogt

## Abstract

Latitudinal diversity gradients (LDGs), typically declining from equator to poles, are a pervasive macroecological pattern, yet their generality and drivers in the ocean microbiome remain widely unresolved. We integrated global-scale metagenomic data with habitat modeling to study marine microbial LDGs across seasons and depths. Surface mixed layer microbiomes exhibited diversity peaks at (sub)tropical latitudes and a poleward decline, whereas mesopelagic communities (200–1,000 m) showed no latitudinal diversity structuring. Taxonomic resolution revealed that the mixed layer LDG was underpinned by Alphaproteobacteria and Cyanobacteriia, while other taxa exhibited distinct or contrasting LDGs. Diversity structuring also varied by seasons and regions, governed by temperature and nutrient availability. Together, these findings highlight that within the ocean microbiome, LDGs are not universal, but lineage-specific ecological strategies and responses to environmental gradients. Our study provides fundamental insights into the structuring of ocean microbiome diversity and lays the foundation for predicting responses to environmental change.

## Introduction

Latitudinal diversity gradients (LDGs) have been recognized since the early 19^th^ century, when Alexander von Humboldt documented a decline in diversity from the equator to the poles^1^. Since then, LDGs have been observed across a wide range of terrestrial and aquatic ecosystems and organismal groups^2–5^, reflecting the combined influence of ecological and evolutionary processes across environmental gradients. Understanding how and why diversity is structured along latitudinal and environmental gradients is thus critical, as it provides insights into the mechanisms shaping diversity and informs predictions of how ecosystems may respond to global change.

In marine planktonic communities, LDGs have been observed for both eukaryotes and prokaryotes, but their generality and underlying drivers remain unresolved. Early molecular surveys of marine bacteria, based on 16S rRNA gene fingerprinting and clone library sequencing approaches, reported a decline in richness (the number of unique taxa observed in a sample) toward higher latitudes, consistent with the traditional LDG concept^6,7^. However, the expansion of global ocean sampling, together with advances in sequencing technologies, has challenged these findings and revealed a more complex and inconsistent picture. For instance, a study using high-throughput 16S rRNA gene amplicon sequencing in the South Pacific Ocean reported a bimodal LDG for prokaryotes, with peaks at mid-latitudes^8^. Mid-latitudinal peaks were further supported by metagenomic sequencing of globally distributed samples^9^, while proposed alternatives include a diversity plateau within tropical and subtropical regions^10^, and even an absence of LDGs altogether^11^. Together, these contrasting observations highlight the uncertainty surrounding LDG patterns in marine prokaryotes, which likely reflects both methodological differences and variations in the spatial and temporal coverage of datasets.

Statistical models represent a powerful framework for overcoming some of the limitations of inferring general patterns based on observational datasets, which are often sparse and unevenly distributed across spatial and temporal scales^12^. Such models provide the means to explore large-scale distributions of microbial plankton species. They assume that species are not dispersal-limited in the open ocean, a trait consistent with the generally wide geographic ranges of prokaryotes^13^, and that they are primarily controlled in their global distribution by rapidly responding to abiotic environmental factors^14^. Ladau et al.^15^ applied such an approach to ribosomal DNA to investigate the seasonal variation of near-surface ocean LDGs. They found that during winter months, global prokaryotic richness peaks at high latitudes, which would represent an inversion of the classical LDG pattern. While this study provided strong indications for a seasonal fluctuation of diversity, it was constrained by a relatively small dataset and methodological heterogeneity.

With the growing availability of databases containing uniformly processed metagenomic data (e.g.,OMDB^16^; SPIRE^17^), we can now strive to assemble datasets with extensive spatiotemporal coverage to yield global models with higher resolution. Metagenomic sequencing provides a broad genomic context, enabling consistent species-level taxonomic profiling from multiple marker genes (e.g., mOTUs^18^) that is independent of primer-based amplification. This makes it particularly well-suited for meta-analyses integrating data from multiple sources, as it minimizes technical variability that hampers the integration of amplicon-based datasets generated with different primers, sequencing regions, or clustering thresholds.

Here, we combined a systematically processed metagenomic dataset with extensive spatial and temporal coverage with habitat models (CEPHALOPOD^12^) to study LDGs of marine prokaryotes across depths, seasons, and taxonomic ranks. Our analyses revealed pronounced LDGs for both Bacteria and Archaea in the mixed layer, which were absent in the mesopelagic layer. Furthermore, we observed a spatial decoupling of bacterial and archaeal richness in the mixed layer of (sub)tropical regions, seasonal modulation of diversity without inversion of the general LDG pattern, and variations in LDGs across taxonomic ranks. By linking global diversity patterns to environmental drivers, we found that marine prokaryotic lineages occupy structured ecological niches and developed a framework to characterize diversity hotspots for different taxonomic groups.

## Results

### Global dataset and study design

To characterize global patterns in oceanic prokaryotic diversity, we assembled a metagenomic dataset based on the Ocean Microbiomics Database (OMDB; https://omdb.microbiomics.io)^16^. For our analyses, we considered samples obtained within the mixed layer, the depth of the surface ocean that is well-mixed, and samples from the mesopelagic layer (200–1,000 m depth) (**Methods**). In total, 2,469 samples from the mixed layer and 211 mesopelagic samples matched our selection criteria (**Data and code availability**). Spatially, the dataset provides comprehensive coverage across all major ocean basins (**Fig. 1**). The majority of samples from the mixed layer were collected from the Pacific Ocean (∼50%), followed by the Atlantic Ocean (∼28%), with additional coverage extending into polar regions (∼4%). Temporally, the dataset represents sampling efforts spanning all seasons and a latitudinal range from -65° to 85°.

**Figure 1.**
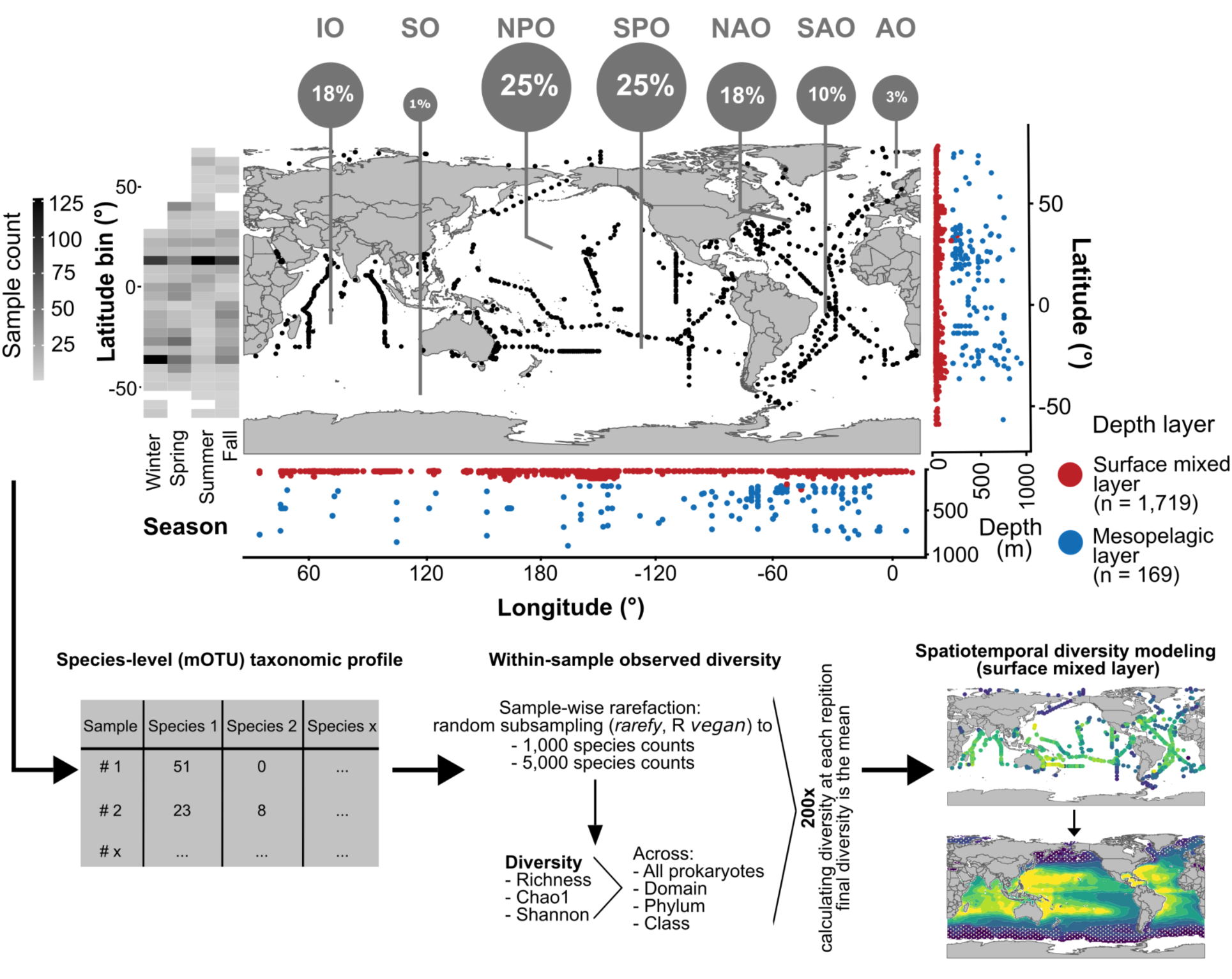
Global dataset of prokaryotic diversity and study design. Global map showing the distribution of metagenomic samples from the surface mixed layer and mesopelagic layer (200–1,000 m). Sample locations and fractions are indicated for the Indian Ocean (IO), Southern Ocean (SO), North Pacific Ocean (NPO), South Pacific Ocean (SPO), North Atlantic Ocean (NAO), South Atlantic Ocean (SAO), and Arctic Ocean (AO). Marginal scatter plots illustrate the vertical sample distribution across the mixed layer (n = 1,719) and mesopelagic layer (n = 169). Sample counts are also shown across aggregated latitude bins (5°) and seasons (left). Seasons were defined as: Northern Hemisphere: winter = Dec–Feb, spring = Mar–May, summer = Jun–Aug, autumn = Sep–Nov; Southern Hemisphere: reversed. Species-level taxonomic profiles were generated and used to calculate within-sample diversity (observed richness, Chao1, Shannon index) across all prokaryotic species, as well as by domain, phylum, and class. For each sample, 200 independent rarefactions were performed, and diversity metrics were calculated as the mean across repetitions. Within-sample diversity metrics were then extrapolated across the mixed layer using habitat modeling.

From these data, we generated taxonomic profiles, resolving species-level metagenomic operational taxonomic units (mOTUs). To account for unequal sampling efforts, we standardized the taxonomic profiles to a fixed number of species observations. Using a rarefaction cutoff of 5,000 species observations, we retained 1,719 samples from the mixed layer and 169 mesopelagic samples (**Methods**). From the rarefied taxonomic profiles, we computed the species-level richness (number of unique species observed per sample), as well as the Shannon^19^ (accounts for both richness and evenness) and Chao1^20^ (estimating unobserved species) diversity indices. These indices were computed across all prokaryotes and at the level of domain, phylum, and class (**Fig. 1**; **Methods**). We subsequently used the resulting species richness after rarefaction to 5,000 species observations as our primary metric for diversity. We first analyzed the diversity pattern emerging from the samples alone, and then expanded our scope using a habitat modeling approach (**Methods**). This allowed us to investigate the annual and seasonal biogeographical structuring of global ocean prokaryotic diversity in the mixed layer.

### Latitudinal diversity gradients are depth-dependent

As a first approach to assess the global distribution of species-level marine prokaryotic diversity, we used within-sample richness to examine LDGs of Bacteria and Archaea in the surface mixed layer and the mesopelagic layer.

In the mixed layer, species-level richness followed a clear LDG, with a ∼3-fold change (FC) increase in richness for both Bacteria and Archaea in tropical and subtropical regions (40° S– 40° N) compared with higher latitudes (Bacteria: FC = 3.1; Archaea: FC = 3.0; both *p* < 0.001, Wilcoxon test; **Fig. 2**). Chao1-estimated richness was likewise higher at (sub)tropical latitudes compared with high latitudes (Bacteria: FC = 2.8, *p* < 0.001; Archaea: FC = 3.1, *p* < 0.001), and Shannon’s diversity index followed the same trend, albeit with smaller fold changes (Bacteria: FC = 1.4, *p* < 0.001; Archaea: FC = 1.6, *p* < 0.001). Prokaryotic richness in the mixed layer was highly correlated with bacterial richness (Bacteria: Pearson’s *r* = 0.998, *p* < 0.001; **Fig. 2a**), consistent with the bacterial dominance of overall prokaryotic richness (96% of total richness; **Supplementary Fig. 1a**). While archaeal richness showed fold changes between low and high latitudes that were similar to those of bacteria, it was only moderately correlated with prokaryotic richness (*r* = 0.34, *p* < 0.001; **Fig. 2**b), suggesting domain-specific biogeographic patterns.

**Figure 2.**
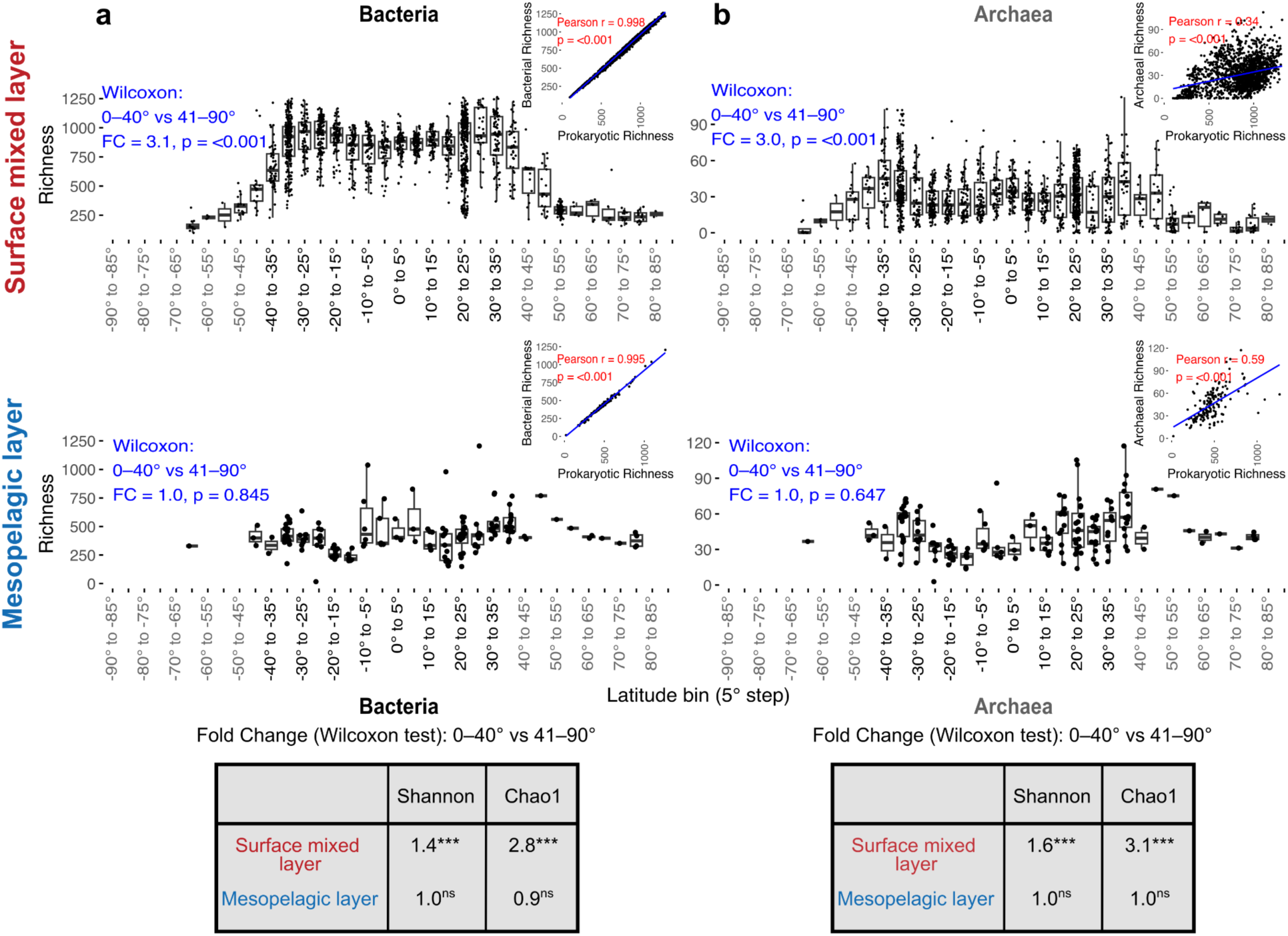
Latitudinal patterns of observed richness in Bacteria and Archaea across ocean depths. Observed within-sample richness for **(a)** Bacteria and **(b)** Archaea in the mixed layer (top panels) and the mesopelagic layer (bottom panels). Statistical comparisons (blue) show fold change (FC) and *p*-value (Wilcoxon test) between (sub)tropical and high-latitude regions (0°–40° vs. 41°–90° absolute latitude). The top-right plots compare the observed richness between all prokaryotes and Bacteria/Archaea. Tables summarize fold changes between 0°–40° and 41°–90° absolute latitude for Shannon index and Chao1 diversity metrics in (a) Bacteria and (b) Archaea (Wilcoxon test; ns: *p* > 0.05, \*\*\**: p* < 0.001).

In contrast to the mixed layer, species-level richness did not differ significantly between low and high latitudes in the mesopelagic layer for either Bacteria or Archaea (Bacteria: FC = 1.0ⁿˢ; Archaea: FC = 0.96ⁿˢ; ns: *p* > 0.05, Wilcoxon test; **Fig. 2**). This absence of a latitudinal gradient was consistent across Chao1 and Shannon indices (FC = 0.9–1.0^ns^ for both domains; **Fig. 2**). To account for sampling differences, we randomly downsampled mixed-layer samples to match the number of mesopelagic samples. The decreased richness for Bacteria and Archaea at high versus low latitudes in the surface mixed layer remained robust even after accounting for sampling differences (**Supplementary Fig. 2**). Thus, our results suggest that mesopelagic diversity does not conform to an LDG characterized by declining richness toward the poles.

The mixed and mesopelagic layers exhibited distinct microbial diversity structures. For Archaea, species-level richness was higher in the mesopelagic (45 ± 19, mean ± SD) than in the surface mixed layer (30 ± 19) (**Supplementary Fig. 1a**). Mixed-layer archaeal richness was largely dominated by members of the class Poseidoniia (27 ± 17), whereas mesopelagic richness was more evenly distributed across Nitrososphaeria (22 ± 11), Nanoarchaeia (2 ± 6), and Poseidoniia (19 ± 12). In contrast, bacterial species-level richness was higher in the mixed layer (799 ± 257) than in the mesopelagic zone (421 ± 158). In the mixed layer, species-level richness was heavily skewed toward a few dominant groups, with species of classes Alphaproteobacteria (430 ± 180) and Cyanobacteriia (106 ± 71) together accounting for 65% of the mean prokaryotic richness (Alphaproteobacteria: 52%, Cyanobacteriia: 13%; **Supplementary Fig. 1b**). Frequently used taxonomic definitions, like the 97% sequence identity clustering of the 16S rRNA gene sequence, result in broader taxonomic resolutions than species level^21^. To test the sensitivity of marine prokaryotic diversity structure to taxonomic resolution, we also calculated richness at the genus level. Genus-level prokaryotic richness in the mixed layer was less strongly dominated by Alphaproteobacteria and Cyanobacteriia, contributing only 31% of the mean prokaryotic richness (Alphaproteobacteria: 29%, Cyanobacteriia: 2%; **Supplementary Fig. 1b**). This indicates that a limited number of genera contributed to the high species-level richness of Alphaproteobacteria and Cyanobacteriia in the mixed layer. In contrast to the species-level richness, genus-level richness was higher in the mesopelagic than in the mixed layer (mesopelagic layer = 237 ± 52, mixed layer = 184 ± 52; **Supplementary Fig. 1c**), consistent with previous reports utilizing taxonomic resolutions broader than species-level^9,22–24^. Differences in the baseline richness across depth layers may also reflect the larger unassigned fraction in the mesopelagic compared to the mixed layer (mesopelagic layer: median = 0.36, mixed layer: median = 0.24; **Supplementary Fig. 1d**). Notably, also at the genus level, there was no significant difference in richness in the mesopelagic between low and high latitude regions (FC = 1.0ⁿˢ; **Supplementary Fig. 1c**). Together, these findings underscore the importance of taxonomic resolution in interpreting microbial diversity patterns across ocean layers.

### Spatial and temporal variations of ocean microbiome diversity in the mixed layer

To extend our analyses and compare mixed-layer archaeal and bacterial LDGs across the global ocean and seasons, we applied a second, habitat modeling-based approach. In this framework, observed diversity was extrapolated in space and time using an ensemble of models trained on environmental variables (**Supplementary Fig. 3**). Global-scale modeled diversity patterns of Bacteria and Archaea were highly consistent across rarefaction cutoffs (Pearson’s *r* ≥ 0.98, all *p* < 0.001; **Supplementary Fig. 4**), indicating that overall results are robust to this methodological choice.

At the domain level, the biogeography of bacterial and archaeal diversity was similar across diversity metrics (Bacteria: richness vs. Shannon, *r* = 0.98, and Chao1, *r* = 0.98; Archaea: richness vs. Shannon index, *r* = 0.87, and Chao1, *r* = 0.99; all *p* < 0.001; **Fig. 3**). Across the global ocean, species-level richness patterns between the two domains showed a moderate correlation (*r* = 0.55, *p* < 0.001; **Supplementary Fig. 5a**). However, within (sub)tropical regions (0°–40° absolute latitude), a decoupling of bacterial and archaeal richness was evident (*r* = - 0.17, *p* < 0.001; **Supplementary Fig. 5b**). This low correspondence was consistent across other diversity metrics (Shannon index: *r* = -0.04, *p* < 0.001; Chao1: *r* = 0.20, *p* < 0.001). Together, these results indicate that (sub)tropical bacterial and archaeal communities follow distinct diversity–latitude relationships, suggesting divergent ecological drivers.

**Figure 3.**
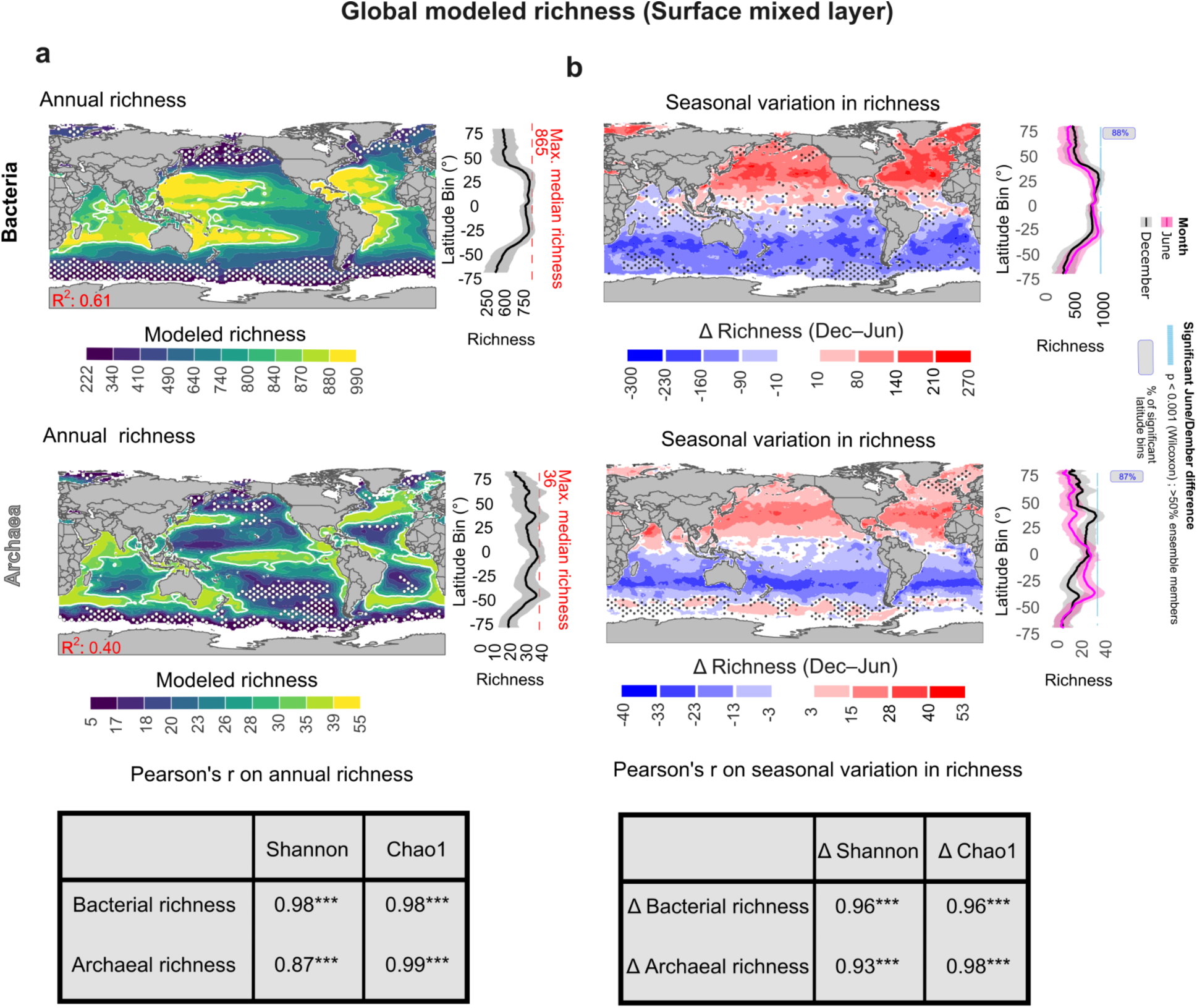
Modeled bacterial and archaeal richness patterns in the mixed layer. **(a)** Global maps of annual richness patterns for Bacteria (upper panel) and Archaea (lower panel), projected using an ensemble-based model. Maps display modeled annual mean richness and areas of uncertainty (white stippling; coefficient of variation > 20% across ensemble members). Model predictive power (R²) is shown in red (lower left). White contours highlight diversity hotspots (regions above the 75^th^ percentile of modeled richness). Latitudinal gradients are shown for the global ocean with shadings indicating model uncertainty (± SD) across ensemble members and longitude. Dashed red lines and annotation (upper right corner) indicate the maximum latitudinal median diversity. **(b)** Seasonal global maps showing the modeled change in richness between December and June for Bacteria (upper) and Archaea (lower). Colors represent the magnitude and direction of change. Black stippling indicates areas of high model disagreement (< 75%) on the direction of richness change across ensemble members. Latitudinal gradients for June and December richness are shown, with uncertainty (± SD) as shading. Horizontal blue labels mark a significant difference between June and December, defined as bins with >50% of ensemble members showing a significant difference (*p* < 0.001; Wilcoxon test); text labels denote the percentage of 1° latitude bins with a significant difference.

The decoupling in annual bacterial and archaeal richness in the mixed layer was also evident in distinct hotspots of diversity, defined as regions above the 75^th^ percentile of predicted richness (**Fig. 3**a). Bacterial hotspots were identified in oligotrophic gyres (∼25° absolute latitude, richness up to 990), while archaeal hotspots occurred in nutrient-rich upwelling zones (e.g., equatorial divergence and ∼40° absolute latitude, richness up to 55). Archaeal richness was markedly lower in oligotrophic regions (richness < 15). Our models show that LDGs vary not only between but also within ocean basins, e.g., bacterial richness dips around the equator in the eastern Pacific, whereas this dip is absent in the western Pacific (**SupplementaryFig. 6a–b**). This indicates that sampling transects in different regions and ocean basins can result in differing LDG shapes.

Our habitat modeling approach also allowed us to identify distinct temporal patterns in the richness of the ocean microbiome. For Bacteria, modeled richness tended to increase during winter months. In the Northern Hemisphere, richness increased by a median of 76 species (12% increase) between June and December (interquartile range (IQR): 21–131), with local increases by up to 279 species (112% increase) at around 25° absolute latitude (**Fig. 3**b). In the Southern Hemisphere, richness declined by a median of 83 species (14% decrease) (IQR: 123 to 46), with a maximum decrease of 165 species (27% decrease) at similar southern latitudes at around - 25°. These seasonal differences were statistically significant across ∼88% of latitudinal bins, defined as those bins where more than 50% of model ensemble members showed significant differences (*p* < 0.001; Wilcoxon test). Archaea showed similar seasonal trends of higher richness in winter than in summer months (**Fig. 3**b). Comparing December to June, local richness maxima were found between 21° and 38° latitude in the Northern Hemisphere, reaching up to 56 species (60% increase), while a minimum was observed between -45° and -30° latitude in the Southern Hemisphere, where richness decreased to 36 species (35% decrease). Statistically significant differences between winter and summer months were observed across 87% of latitudinal bins. Thus, our results indicate an increase in richness for Bacteria and Archaea in winter compared to summer months, whereas previously reported global diversity maxima at high latitudes during winter^15^ are not supported by our data.

While we cannot evaluate the fidelity of the modeled seasonal changes in the open ocean due to the lack of time-series observations, our modeled seasonal changes are reflected in coastal marine systems. Concretely, we compared the prokaryotic difference in richness (ΔRichness_December-June_) for seven European coastal time series stations with the corresponding modeled latitudinal bins (**Supplementary Fig. 7**). All coastal time series and open ocean models showed higher richness in December than in June, with similar median increases (Modeled: 53.9%, Time series: 56.9%; **Supplementary Fig. 7**). This suggests that prokaryotic richness is higher in winter than in summer months, across both open ocean and coastal environments.

### Latitudinal diversity gradients vary across taxonomic groups

To gain deeper insights into the structure of bacterial and archaeal richness across the global ocean mixed layer, we modeled species-level richness per class and derived their LDGs (**Supplementary Fig. 10**). We first compared annual LDGs across classes to assess whether they are uniform or vary. We then assessed whether similarities in richness distributions align with phylogenetic relationships, and finally investigated how seasonal variability contributes to diversity dynamics across taxonomic groups.

A comparative assessment of annual LDGs across classes revealed four main groups of diversity gradients (**Fig. 4**a). The bacterial LDG was highly similar to that of the two richest classes, Alphaproteobacteria (*r* = 0.998, *p* < 0.001) and Cyanobacteriia (*r* = 0.970, *p* < 0.001) (Group 2; **Fig. 4**a), highlighting that the overall prokaryotic and bacterial LDG is primarily shaped by species of these two classes. Similarly, clustering in group 4 reveals that the archaeal LDG is highly aligned with the LDG of class Poseidoniia (*r* = 0.976, *p* < 0.001). Despite variation in gradient shape, most taxonomic groups showed lower diversity at temperate and polar latitudes compared to (sub)tropical zones. Notable exceptions included group 1, with Archaea of the class Nitrososphaeria and Bacteria of the class Fibrobacteria, both showing increasing diversity toward high latitudes (Nitrososphaeria: *r* = 0.71, *p* < 0.001; Fibrobacteria: *r* = 0.79, *p* < 0.001). Together, masked by species-rich taxonomic groups, particularly Alphaproteobacteria and Cyanobacteriia, a variety of LDGs exist in the surface mixed layer.

**Figure 4.**
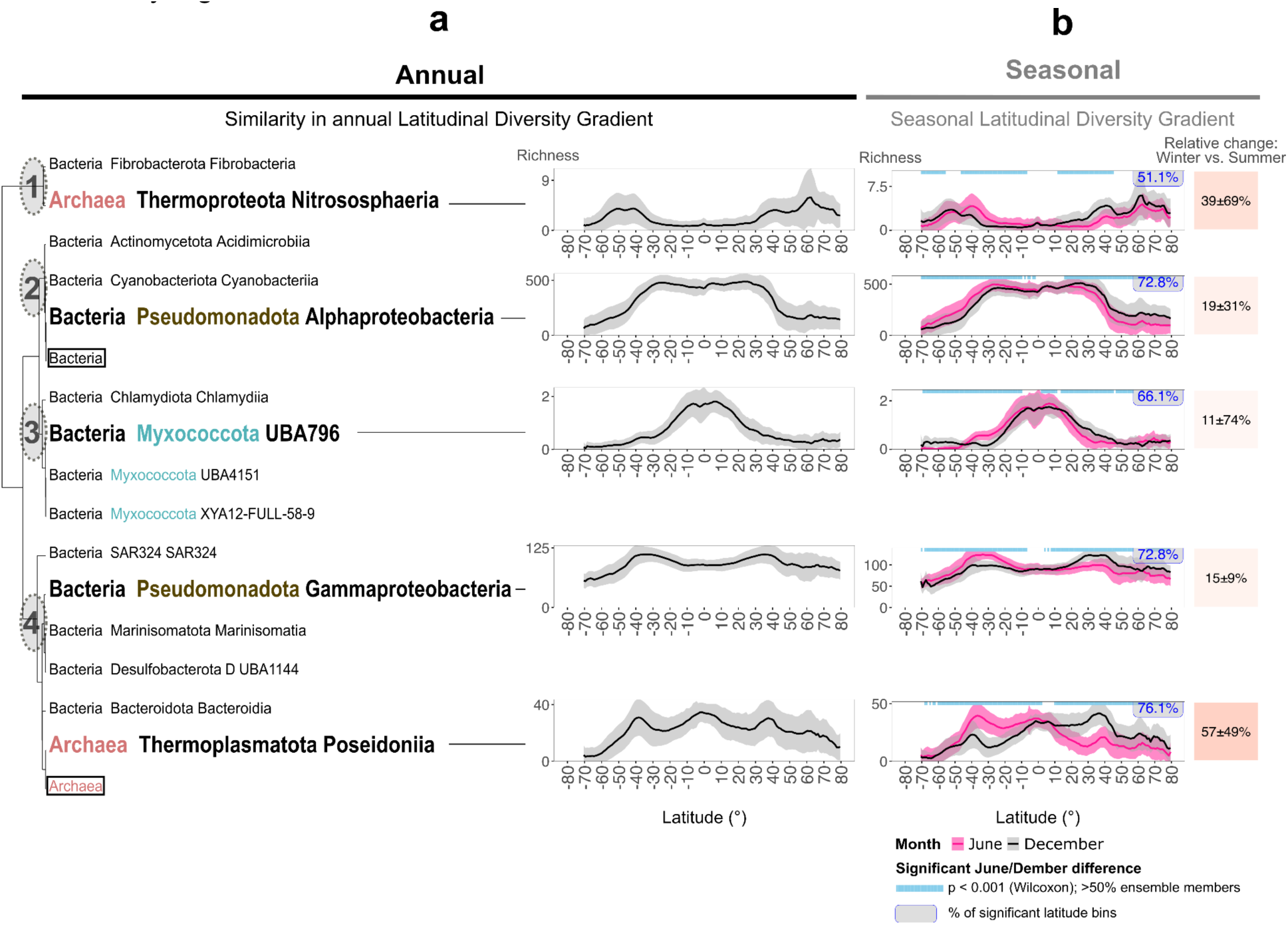
Latitudinal diversity gradients across taxonomic ranks and seasonality. **(a)** Dendrogram represents the similarity in model-based annual latitudinal richness patterns across taxonomic groups. Mean richness per 1° latitude bin was correlated between taxonomic ranks (Pearson correlation; Hierarchical clustering method “ward.D2”). Four distinct groups of latitudinal diversity gradients (LDGs) were identified, henceforth called groups. For each group, the representative taxonomic rank with the highest average annual richness in the mixed layer is shown, plus the archaeal class with the highest average richness (Poseidoniia). Solid lines represent modeled mean richness per 1° latitude bin, with shaded areas indicating the standard deviation across model ensemble members and longitude. **(b)** Left: Modeled mean latitudinal richness for June (pink) and December (gray). Horizontal blue labels mark a significant difference between June and December, defined as bins where > 50% of the ensemble members showed a significant difference (*p* < 0.001; Wilcoxon test); text labels indicate the percentage of 1° latitude bins with a significant difference. Right: The relative richness change was calculated as the mean richness in winter (NH: December; SH: June) relative to summer (NH: June; SH: December) ± SD across all latitude bins.

To investigate the relationship between annual LDGs among taxonomic groups (**Fig. 4**a) and their phylogenetic distances, we compared latitudinal richness distributions to patristic distances. Alignment of both dendrograms revealed disparities (entanglement score = 0.6; **Supplementary Fig. 8**). For example, although the classes Alphaproteobacteria and Gammaproteobacteria belong to the same phylum, their LDGs differed considerably (*r* = 0.71, *p* < 0.001) and did not group together (Alphaproteobacteria: Group 2, Gammaproteobacteria: Group 4; **Fig. 4**a). Therefore, observed (dis)-similarities in the shape of latitudinal richness distributions do not reflect phylogenetic distance.

Each of the four identified LDG groups also exhibited a significant seasonal change in richness in more than half of the latitude bins, thereby contributing substantially to the seasonal changes of the whole community (**Fig. 4**b). Here, the seasonal change was calculated as the ratio of mean winter month richness (December in the Northern Hemisphere, June in the Southern Hemisphere) to mean summer richness (June in the Northern Hemisphere, December in the Southern Hemisphere), averaged across all latitude bins. Seasonal changes in richness consistently increased during winter months compared to summer months, across taxonomic groups. The magnitude of seasonal change in relative richness varied among representative classes, ranging from 57 ± 49% in Poseidoniia to 11 ± 74% in UBA796 (mean ± SD). Notably, the LDG of class UBA796, which showed the highest richness around the equator where seasonality is limited, exhibited minimal seasonal change. These results indicate that higher richness in winter is a widespread trend across bacterial and archaeal taxonomic groups. However, the magnitude of seasonal change varied among classes and across latitude, as indicated by high standard deviations.

### Divergent environmental drivers and global diversity hotspots across taxonomic groups

Having identified four main groups of annual LDGs across taxonomic ranks in the mixed layer (**Fig. 4**), we sought to characterize their key environmental drivers and contextualize their diversity hotspots (**Fig. 5**). To disentangle the environmental drivers of prokaryotic diversity, we first assessed correlations among climatological features, which grouped into clusters representing temperature, oxygen, macronutrients, and carbon dynamics (**Supplementary Fig. 9**). Given the strong covariation among variables in the surface ocean, predictor importance reflects dominant environmental regimes rather than individual variables.

**Figure 5.**
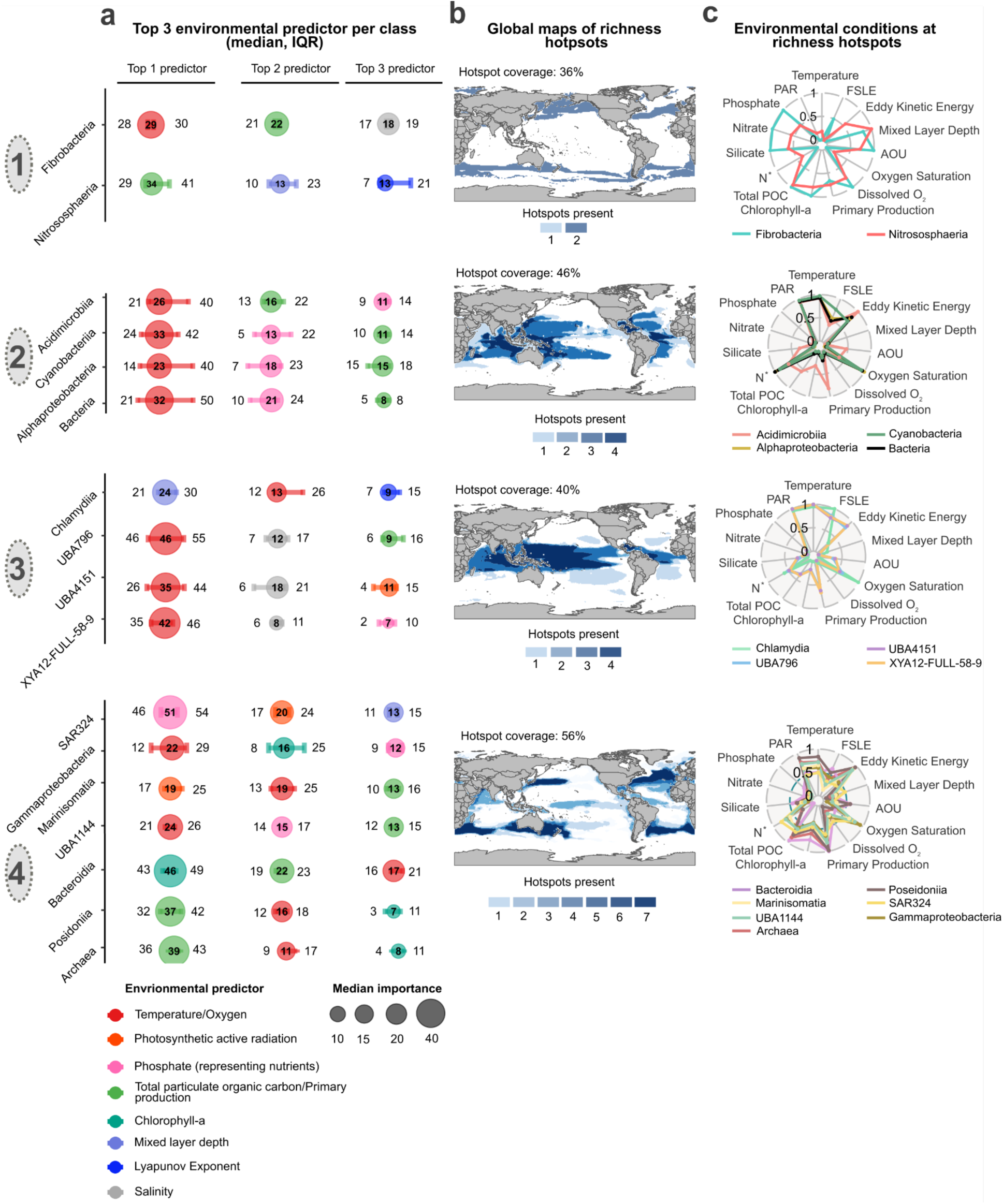
Richness hotspots and environmental drivers in the surface mixed layer across taxonomic ranks. **(a)** Bubble plots show the top three environmental drivers of richness for each prokaryotic class and domain (color-coded). Bubble size represents the median variance explained across ensemble members, while horizontal bars show the interquartile range across ensemble members (different algorithms and bootstrap iterations). **(b)** Consensus hotspot maps show, for each LDG group (Fig. 4), the number of classes with richness hotspots at a location (annual modeled richness > 75^th^ percentile) and the fraction of global ocean area covered by that group’s hotspots (spatial resolution 1° x 1°). **(c)** Spider plots depict scaled environmental characteristics (annual mean) of each class across global diversity hotspots. Environmental variables were scaled to the range 0-1 using min-max normalization (R scales::rescale function), to allow direct comparisons. For example, a temperature value of 1 indicates that the richness hotspot for that class has the highest median annual temperature compared to all other lineage-specific richness hotspots. Annual mean environmental parameters quantified are: temperature, photosynthetically active radiation (PAR), phosphate, nitrate, silicate, N* (excess nitrate relative to phosphate), total particulate organic carbon (POC), chlorophyll-a, primary production, dissolved oxygen (O₂), oxygen saturation, apparent oxygen utilization (AOU), mixed layer depth, eddy kinetic energy, and Lyapunov exponent (FSLE). Lines are color-coded by class.

Across taxonomic groups, temperature consistently emerged as the most important driver of richness (9 out of 15 classes), particularly for classes within group 2 (e.g., Acidimicrobiia: 28%, IQR: 21–40%, median and interquartile range) and group 3 (e.g., XYA12-FULL-59-9: 42%, IQR: 35–46%) (**Fig. 5**a). Total particulate organic carbon (POC) levels, correlated with primary production (*r* = 0.84, *p* < 0.001; **Supplementary Fig. 9**), emerged in 10 out of 15 classes as a secondary or tertiary driver and even acted as the strongest predictor in some cases (e.g., group 4: POC for Poseidoniia: 37%, IQR: 32–42%). Beyond that, phosphate, covarying with other inorganic macronutrients, was identified as a secondary or tertiary driver in 7 classes (e.g., in all classes of group 2, ranging from Acidimicrobiia: 11%, IQR: 9–14% to Alphaproteobacteria: 18%, IQR: 7–23%). These results highlight that, while temperature exerts a primary influence on richness, lineages exhibit distinct sensitivities to macronutrient and carbon gradients, pointing to divergent ecological strategies.

The environmental characteristics and spatial extent of diversity hotspots (area of > 75^th^ percentile modeled richness; **Supplementary Fig. 10**) are complementary to the dominant environmental features identified for each LDG group (**Fig. 4**a). Group 1 classes (Nitrososphaeria and Fibrobacteria) had richness hotspots in polar and subpolar waters, representing nutrient- and carbon-rich regimes, and their richness hotspots occupied ∼36% of the global mixed layer (**Fig. 5**b). In contrast, group 2 clades (classes Acidimicrobiia, Cyanobacteriia, Alphaproteobacteria, as well as the overall domain Bacteria) were richest in oligotrophic (sub)tropical waters, with highest temperature and lowest phosphate concentrations across all groups (**Fig. 5**c). Their richness hotspots spanned ∼42% of the global ocean but were absent from more nutrient-rich equatorial upwelling zones (**Fig. 5**b). Group 3 (class Chlamydiia and three Myxococcota classes: UA796, UBA4151, XYA12-FULL-59-9) were geographically constrained to richness hotspots around the equator (∼40% of the global mixed layer), characterized by high photosynthetically active radiation (PAR) and a shallow mixed layer (**Fig. 5**c). Finally, group 4 (classes SAR324, Gammaproteobacteria, Marinisomatia, UBA1144, Bacteroidia, Poseidoniia, as well as the overall domain Archaea) exhibited the broadest extent of richness hotspots (∼56% of the global mixed layer ocean), concentrated at mid-latitudes outside of the oligotrophic gyre regions, under elevated nutrient concentrations and POC levels (**Fig. 5**c). Taken together, richness hotspots and ultimately LDGs were shaped by lineage-specific associations with environmental gradients, driven primarily by temperature and secondarily by macronutrient and organic carbon availability.

## Discussion

Here, we show that the latitudinal structuring of species-level diversity in the ocean microbiome is depth-dependent and shaped by lineage-specific biogeographic dynamics. Across the surface mixed layer, the prokaryotic LDG is mainly structured by classes with diversity peaks in subtropical gyres, while other taxonomic groups show diversity maxima at equatorial or higher latitudes. These contrasting patterns underscore taxon-specific responses to environmental drivers, including temperature, nutrient, and organic carbon concentrations and phytoplankton productivity. Thus, our findings suggest that the diversity of the ocean microbiome does not universally conform to the classical LDG perspective but instead is shaped by the interaction between distinct ecological strategies and environmental gradients that result in taxon-specific diversity patterns.

The depth-dependent nature of LDGs suggests that microbial diversity is shaped by fundamentally different drivers in the surface mixed layer and mesopelagic environments. In the mixed layer, environmental conditions, such as temperature, light, and nutrient availability, fluctuate over daily to seasonal timescales^4^ (**Supplementary Fig. 3**). These variations create heterogeneous ecological opportunities, driving microbial responses that underpin the structuring of diversity over space and time. As shown by Villarino et al.^25^, community composition in surface prokaryotic communities changes in response to environmental differences and exhibits more pronounced variability in surface waters, whereas communities in the mesopelagic layer are more similar and influenced by longer-term processes such as water-mass circulation and residence time^26^. Arctic time-series observations further show that while surface microbial communities shift markedly with light availability and phytoplankton bloom dynamics, deeper mesopelagic assemblages remain comparatively stable across seasons^26^. The mesopelagic layer is characterized by relatively stable physical conditions, such as temperature and oxygen, across latitudes, yet it experiences strong temporal variations in the input of organic matter. Although temporal variability remains pronounced for such biological parameters, the lack of a strong latitudinal structuring likely contributes to the absence of a latitudinal diversity gradient in the mesopelagic. However, given the lower sampling coverage of the mesopelagic layer, further sampling and investigation are needed to improve our understanding of diversity structuring and the processes shaping microbial communities below the ocean’s surface.

The pronounced LDGs in the surface mixed layer are structured over both spatial and temporal scales. Our results do not support a seasonal inversion of the bacterial LDG with diversity peaks in (sub)polar regions in winter months, as previously hypothesized^15^. Instead, our results suggest that the classical LDG pattern (poleward decrease in diversity) holds in the surface mixed layer throughout the year, but its shape is modulated by seasonal dynamics: In agreement with observations from time-series studies^24,27–29^, our model projects an increase in bacterial and archaeal diversity in winter relative to summer months. A major factor that likely contributes to seasonal variations is the deepening of the surface mixed layer in winter, which leads to vertical mixing of previously stratified communities, thereby increasing species richness and evenness^28,29^. By capturing spatiotemporal dynamics, our findings indicate that previously reported deviations in LDG shape may be partially explained by seasonal and spatial variability. Our model can serve as a reference to incorporate temporal dimensions into the monitoring of global ocean microbiome diversity.

We identified divergent LDGs at the class level and a domain-specific decoupling of richness between Bacteria and Archaea in the (sub)tropical mixed layer. Although global LDGs have been reported separately for both domains^8,10^, their spatial decoupling has not previously been highlighted. This decoupling was largely underpinned by Cyanobacteriia and Alphaproteobacteria species, which dominate the bacterial diversity signal and exhibit richness hotspots in oligotrophic gyres, consistent with their adaptations to nutrient-scarce, low-productivity environments^30,31^. These groups display diverse ecological strategies, encompassing both photoautotrophic and heterotrophic energy acquisition, which confer varying dependencies on light availability, organic substrate utilization, temperature tolerance, and resource efficiency^30,32,33^. Such functional versatility promotes a high degree of niche partitioning^34,35^ and helps explain the elevated bacterial richness observed in these regions^36^, ultimately driving LDG peaks at intermediate latitudes. Archaeal richness, meanwhile, is largely structured by Poseidoniia species, which have been linked to productive waters with the availability of phytoplankton-derived dissolved organic matter^37,38^. Beyond these dominant groups, distinct LDG patterns were also evident in other classes. For example, many Bacteroidia species respond to organic matter available in upwelling systems^39^, which can explain their LDG shape that closely follows the gradient of primary production (**Supplementary Figs. 3 and 10**). Similarly, Gammaproteobacteria show richness hotspots outside of oligotrophic gyres, consistent with their classification as copiotrophs that preferentially occupy nutrient-rich environments^30,40^. The archaeal class Nitrososphaeria (phylum Thermoproteota) displayed elevated richness levels at higher latitudes, in alignment with reports of their high prevalence in surface waters around Antarctica^41^. Their rare abilities to yield energy from ammonia oxidation and urea degradation^42^ and to withstand freeze–thaw cycles^43^ likely enable their unusual LDG. Taken together, these examples underscore that surface mixed-layer LDGs are not universal features but arise from taxon-specific ecological and evolutionary adaptations to distinct environmental regimes.

Despite such taxon-specific differences, several broader patterns emerge, indicating a degree of generality in the structuring of diversity and its environmental drivers. Most classes share a characteristic decline in diversity toward (sub)-polar regions (**Fig. 4**), with temperature identified as a first-order driver (**Fig. 5**), consistent with macroecological principles linking temperature to biodiversity patterns^4,5,45^. Notably, this shape is consistent with a bimodal structuring of marine diversity, where richness peaks at mid-latitudes and a dip occurs near the equator^8,9,46^. Such bimodality has been hypothesized to arise from processes acting at the thermal boundaries of the tropics, where adaptation to changing temperature regimes may promote diversification, while permanent oceanographic features such as the equatorial upwelling could act as physiological or ecological filters limiting persistence and richness^8,46^. Temperature influences metabolic rates, energy flux, and ecosystem productivity, which collectively constrain microbial diversity across latitudes^45,47^. Alongside this, secondary and tertiary drivers, including the concentration of particulate organic carbon and phosphate, correlated with other macronutrients, were shared across a majority of classes. The observed variations among taxa highlight the limits of universal environmental features, showing instead that diversity patterns emerge from taxon-specific adaptations to distinct combinations of temperature, light, nutrients, and carbon dynamics.

While our global model projections enabled the disentangling of taxon- and depth-specific LDGs within the ocean microbiome and their associated environmental drivers, several limitations remain. A major source of uncertainty stems from the uneven spatial and temporal sampling coverage across the global ocean. This is particularly evident at high latitudes and in the Southern Ocean, where data scarcity restricts our ability to fully capture prokaryotic diversity dynamics and highlights the need for targeted sampling efforts. Additionally, due to the limited spatial and temporal coverage of available global environmental datasets, our models are most robust for open-ocean pelagic environments and less able to resolve highly dynamic coastal regions. Furthermore, while we identified strong associations between environmental variables and diversity patterns, the high degree of covariation among environmental features highlights the challenge of disentangling individual drivers. These relationships should therefore be further explored through experimental and mechanistic studies to better understand the causal mechanisms underlying observed diversity patterns.

Our results demonstrate that global patterns of ocean prokaryotic diversity are shaped by species of a few dominant classes whose richness aligns with distinct environmental drivers, primarily temperature, nutrients, and carbon-related variables. These environmental dependencies reflect fundamental ecological strategies, with some clades thriving in nutrient-poor, stratified waters and others linked to nutrient-rich, dynamic regions. Consequently, future changes in ocean stratification, nutrient availability, and primary production are expected to impact microbial diversity and ecosystem functions unevenly across taxonomic groups. Given the central role of marine prokaryotes in sustaining biogeochemical cycles, understanding their diversity is critical for assessing the resilience of ocean ecosystems under increasing environmental stress. These findings highlight the need for taxon-resolved, environmentally explicit models to accurately forecast microbiome dynamics and their biogeochemical consequences in a changing ocean.

## Methods

### Metagenomic sample selection and generation of taxonomic profiles

Metagenomic data were obtained from the Ocean Microbiomics Database version 2 (OMDB)^16^, which compiles marine metagenomic datasets from publicly available studies and BioProjects registered in the European Nucleotide Archive (ENA). Only metagenomic data used in published studies or shared via personal communication were included. Raw sequencing reads were processed following the procedure described by Doytchinov and Dimov^48^, including adapter trimming, quality filtering, and removal of contaminant sequences. Read sets from large datasets, such as Tara Oceans, were normalized to optimize assembly. Metagenomes were individually assembled using metaSPAdes, and scaffolds (≥1 kbp) were retained for downstream analysis. Metagenome-assembled genomes (MAGs) were reconstructed by mapping quality-filtered reads to assembled scaffolds, estimating coverage across samples, and binning contigs using MetaBAT 2^49^. MAG quality was assessed with CheckM^50^ and anvi’o^51^, retaining genomes with high completeness and low contamination. Species-level clustering was performed using dRep based on average nucleotide identity, and genomes were taxonomically annotated with GTDB-Tk (GTDB R220). Taxonomic profiles were generated using mOTUs 4 default parameters^52^. The final dataset, including genome annotations and metadata, is available in OMDB (https://omdb.microbiomics.io/).

For the diversity analysis of marine communities, we excluded samples from coastal environments (within 20 km of the shoreline) to minimize the confounding effects of eutrophic and dynamic coastal processes and further excluded samples from the enclosed basins Baltic and Mediterranean Sea. Sensitivity analyses were conducted by iteratively excluding samples based on their distance from the coast to assess model stability (**Supplementary Fig. 11**). Additionally, samples collected in response to specific events (e.g., algal blooms, pollution events) and those with a minimum filter size > 0.8 µm were removed to filter out samples that exclude the free-living prokaryotic community, retaining samples with lower filter size thresholds between 0.2–0.8 μm. For surface mixed layer analyses, we retained samples from within the mixed layer depth at each location, with depth estimates derived from the World Ocean Atlas 2018 (WOA18)^53^. Samples collected from depths between 200 and 1,000 m, and located below the mixed layer, were classified as mesopelagic.

### Diversity indices

To account for inter-sample differences in sampling effort and sequencing depth, mOTUs counts were rarefied to equal counts (*rrarefy* function, *vegan* R package). Rarefaction was conducted with two different sample sizes (1,000 and 5,000 mOTU counts), considering only samples with mOTU counts above the respective threshold. Rarefying was conducted 200 times per sample. At each rarefaction repetition, the rarefied mOTUs table was subsetted to species belonging to the same clade (domain, phylum, class), and diversity metrics (richness, Shannon index, Chao1) were calculated. Final diversity values per sample and clade represent the mean of the 200 repetitions.

### Environmental features

Environmental parameters used to model prokaryotic diversity indices capture physical, chemical, and biological characteristics of the marine environment, serving as key environmental features of prokaryotic diversity^6,8,9^. Biological variables include chlorophyll-a concentration (mg m⁻³), a proxy for phytoplankton biomass, and primary production (mg C m⁻² day⁻¹), both sourced from GIMS climatologies^54,55^. Physical parameters encompass sea surface temperature (°C) from the World Ocean Atlas 2018^53^ and photosynthetic active radiation (Ein m⁻² day⁻¹) from the Global Marine Information System^55^ as well as oceanographic dynamics metrics such as eddy kinetic energy (cm² s⁻²)^56^ and the Lyapunov exponent (days⁻¹; obtained from AVISO^57^). Chemical environmental features include nitrate, phosphate, and silicate concentrations (μmol kg^-1^) from the World Ocean Atlas 2018, along with derived stoichiometric indices such as N-star ([Nitrate] - 16*[Phosphate]) and Si-star ([Silicate] - [Nitrate]) to assess nutrient limitation patterns. Oxygen-related variables, including concentration (μmol kg^-1^), fractional saturation (%), and apparent oxygen utilization (μmol kg^-1^), are also sourced from the World Ocean Atlas 2018^53^. Additionally, total particulate organic carbon (mg C m⁻³), an indicator of organic matter availability, was derived from Stramski et al.^58^. All climatologies have a monthly spatial resolution of 1° x 1° longitude and latitude, and annual means alongside seasonal latitudinal variations are visualized in **Supplementary Fig. 3**. Although this monthly resolution does not capture finer temporal dynamics of marine microbial communities, it has proved sufficient to represent major seasonal trends^59,60^ and allows the use of robust climatology datasets that enhance the signal-to-noise ratio in downstream analyses.

### Habitat modeling framework

To model microbial diversity patterns, we employed CEPHALOPOD (Comprehensive Ensemble Pipeline for Habitat modelling Across Large-scale Ocean Pelagic Observation Datasets)^12^, a state-of-the-art, standardized, and highly automated habitat modeling framework designed to produce intercomparable outputs across scarce observation datasets, and observation types, such as metagenomics. CEPHALOPOD follows an ensemble modeling approach, including regression and machine learning algorithms, evaluated by an explicit traffic-light quality check system that ensures robust predictions of the biological target(s), which in this study were the diversity metrics richness, Shannon index, and Chao1. Here, we briefly recall the main steps of CEPHALOPOD and refer to Schickele et al.^12^ for detailed methods and quality check benchmarks.

First, CEPHALOPOD matches metagenomic observations to the resolution of the environmental features to ensure spatial and temporal consistency between the biological target(s) and the environmental conditions in which they were observed. Second, features showing negligible association with the target variables are discarded. Then, CEPHALOPOD only retains the most informative features among groups of intercorrelated ones. Finally, CEPHALOPOD performs a recursive feature elimination procedure, only retaining the minimal, non-redundant set of environmental features best explaining the variance in the target. At this stage, we retain a robust, non-collinear, non-redundant, and parsimonious set of environmental features to best model diversity metrics.

The selected features were then used in a standardized ensemble habitat modeling approach, balanced across regression (GLM; GAM), tree-based (RF, XGBoost), and machine learning algorithms (SVM, MLP). To mitigate the effects of spatial autocorrelation, which can bias algorithm performance evaluation, and to optimize both algorithm training and hyperparameter tuning, CEPHALOPOD implements a spatial block cross-validation procedure. This provides an effective balance between spatial independence and sufficient sample size for algorithm calibration and evaluation. Algorithm skill is then assessed by comparing predicted and observed target values across the evaluation folds, while the variance explained by each feature is quantified to assess model interpretability. Both performance and feature importance are associated with quality checks (i.e., R^2^ > 0.25 and top 3 features > 50 % variance explained). To propagate uncertainty to the projections, CEPHALOPOD generates bootstrap replicates across all algorithms passing the above-mentioned quality checks. Finally, the variability in the projections, across bootstraps and algorithms, is associated with a final quality check (i.e., normalized standard deviation < 0.5). This ensures that only robust and interpretable algorithm outputs are considered in subsequent analyses.

### Selection of modeled taxonomic groups

Annual and seasonal diversity distribution in the ocean mixed layer was modeled across taxonomic levels (**Fig. 4**). We restricted this to domains, phyla, and classes that could be reliably modeled (achieving an R² ≥ 0.25). If the class-level annual LDG was highly correlated with that of its corresponding phylum (*r* > 0.9), we only showed the finer taxonomic resolution (class level) to avoid redundancy. For each modeled phylum with R² ≥ 0.25 (n = 11 phyla), the annual LDG of at least one corresponding class was highly correlated with the annual LDG of its corresponding phylum.

### Statistical testing of seasonal variation in diversity

To assess seasonal variations in marine prokaryote diversity along latitudinal gradients, richness data for December (winter) and June (summer) were extracted from ensemble mean outputs of diversity models. These data were grouped into 1° latitude bins, and median richness values were calculated for each bin and month. Variability around the median was quantified as ±1 standard deviation (SD). Statistical differences in richness between December and June across latitude bins were evaluated using the Wilcoxon test, implemented via the wilcox.test() function in R. Latitude bins with significant differences (*p* < 0.05) were identified, and their significance was visually distinguished by adjusting the transparency (alpha) of the plotted lines. Results were visualized using the R *ggplot2* package, where solid lines represented median richness values, and shaded ribbons depicted variability (± SD). To assess the sensitivity of statistical significance to bin size choices, the analysis was repeated using different latitude bin sizes (1°, 2°, 5°, and 10°). The proportion of statistically significant bins (*p* < 0.001) remained consistently high across all bin sizes, ranging from 93.8% to 96.1%, indicating that bin size had minimal influence on the overall significance of results. This suggests that the observed seasonal differences in prokaryote richness across latitudinal gradients are robust to bin size selection. To account for model uncertainty, we evaluated seasonal differences in richness across individual ensemble members, defined by unique combinations of bootstrap resampling and algorithm choices. Instead of relying solely on the ensemble mean, Wilcoxon tests were performed separately for each ensemble member within 1° latitude bins, allowing a more comprehensive assessment of variability. The proportion of ensemble members displaying significant seasonal differences (*p* < 0.001) was then calculated to quantify the robustness of observed patterns across models. This approach ensures that conclusions about seasonal richness shifts are not overly influenced by any single modeling choice but instead reflect consistent patterns across a diverse set of predictive algorithms and resampling strategies. Visualization of results included both median richness estimates and variability across ensemble members, with shaded confidence regions representing the range of model outputs.

### European time series studies - Seasonality

All data used for the seasonal analysis of European time-series were obtained from the MicrobeAtlas^61^ download section (**Supplementary Table 1**). Richness was calculated from operational taxonomic unit (OTU) read counts (97% sequence identity) using the *rrarefy* function of the R *vegan* package. Per sample, the OTU read counts were rarefied to 3,000 observations with 20 repetitions. Final richness values were obtained by calculating the median richness of the 20 rarefying rounds. Within each study, we summarized richness by sampling month (median) and then calculated the relative change between December and January (**Supplementary Fig. 7**).

### Declaration of generative AI technology in the writing process

During the preparation of this work, the authors used ChatGPT (OpenAI) in order to improve language and readability. After using this tool/service, the authors reviewed and edited the content as needed and take full responsibility for the content of the published article.

## Data and code availability

All contextual information for the metagenomic samples, including associated metadata and calculated diversity metrics for each sample, together with the code used to conduct the analyses presented in this work, are available at https://github.com/grp-bork/LDG_Eriksson_Schiller_2025. The code used to implement the habitat modeling pipeline is openly available at https://github.com/alexschickele/CEPHALOPOD/.

## Author contributions

D.E., J.S., S.S., N.G. M.V. and P.B. were responsible for the conceptualization of this study, further refined by T.P., L.J.U., A.M. and M.K.; Supervision by N.G., P.B., S.S. and M.V.; Data curation was conducted by S.M.V., H.J.R., and C.C.; Analysis and Investigation was led by J.S. (diversity metrics) and D.E. (spatiotemporal modeling), with contributions from E.F. and A.M. (European time series and phylogenetic analysis); Funding was acquired by P.B, S.S., M.V. and N.G; Bioinformatic methodologies applied or adjusted by D.E., A.S. (CEPHALOPOD Model), J.S. (taxonomic resolution) and S.M.V. oversaw resources; Visualizations by D.E. and J.S.; Writing by D.E. J.S. N.G. S.S. and M.V., and the feedback of all other authors.

## Declaration of interests

The authors declare no competing interests.

## Acknowledgements

This work was supported by the Swiss National Science Foundation (SNSF), grant no. 205320_215395. This project has also received funding from the European Union’s Horizon 2020 Research and Innovation Programme under grant agreement no. 862923 (AtlantECO) and no. 101059915 (BIOcean5D). This output reflects only the author’s view, and the European Union cannot be held responsible for any use that may be made of the information contained therein. S.M.V. acknowledges funding from the Human Frontier Science Program (HFSP) through a postdoctoral fellowship [LT0050/2023-L]. We would also like to acknowledge the ETH IT services and HPC facilities for granting access to the EULER high performance cluster. This work was supported by the EMBL IT Services HPC resources (DOI 10.5281/zenodo.12785829). The authors thank Janko Tackmann and Christian von Mering for helpful discussions around the coastal seasonal European time series analyses.

## Notes

https://github.com/grp-bork/LDG_Eriksson_Schiller_2025

https://github.com/alexschickele/CEPHALOPOD/

https://omdb.microbiomics.io

### Competing Interest Statement

The authors have declared no competing interest.

### Summary of Updates

The abstract has been shortened, and the plot subtitles for Mesopelagic and Surface Mixed Layer in Supplementary Figure 1c have been switched.

## References

1. Von Humboldt, A. & Bonpland, A. (Levrault, Schoell et Cie, Paris, 1807).

2. Economo, E. P. et al. Evolution of the latitudinal diversity gradient in the hyperdiverse ant genus *Pheidole*. Glob. Ecol. Biogeogr. 28, 456–470 (2019).

3. Miller, E. C. & Román-Palacios, C. Evolutionary time best explains the latitudinal diversity gradient of living freshwater fish diversity. Glob. Ecol. Biogeogr. 30, 749–763 (2021).

4. Righetti, D., Vogt, M., Gruber, N., Psomas, A. & Zimmermann, N. E. Global pattern of phytoplankton diversity driven by temperature and environmental variability. Sci. Adv. 5, eaau6253 (2019).

5. Benedetti, F., Gruber, N. & Vogt, M. Global gradients in species richness of marine plankton functional groups. J. Plankton Res. 45, 832–852 (2023).

6. Fuhrman, J. A. et al. A latitudinal diversity gradient in planktonic marine bacteria. Proc. Natl. Acad. Sci. 105, 7774–7778 (2008).

7. Pommier, T. et al. Global patterns of diversity and community structure in marine bacterioplankton. Mol. Ecol. 16, 867–880 (2007).

8. Raes, E. J. et al. Oceanographic boundaries constrain microbial diversity gradients in the South Pacific Ocean. Proc. Natl. Acad. Sci. 115, (2018).

9. Sunagawa, S. et al. Structure and function of the global ocean microbiome. Science 348, 1261359 (2015).

10. Ibarbalz, F. M. et al. Global Trends in Marine Plankton Diversity across Kingdoms of Life. Cell 179, 1084–1097.e21 (2019).

11. Moss, J. A., Henriksson, N. L., Pakulski, J. D., Snyder, R. A. & Jeffrey, W. H. Oceanic Microplankton Do Not Adhere to the Latitudinal Diversity Gradient. Microb. Ecol. 79, 511–515 (2020).

12. Schickele, A. et al. CEPHALOPOD, a package to standardize marine habitat-modelling practices and enhance inter-comparability across biological observations. Methods Ecol. Evol. 2041–210X.70040 (2025) doi:10.1111/2041-210X.70040.

13. Whittaker, K. A. & Rynearson, T. A. Evidence for environmental and ecological selection in a microbe with no geographic limits to gene flow. Proc. Natl. Acad. Sci. 114, 2651–2656 (2017).

14. Bunse, C. & Pinhassi, J. Marine Bacterioplankton Seasonal Succession Dynamics. Trends Microbiol. 25, 494–505 (2017).

15. Ladau, J. et al. Global marine bacterial diversity peaks at high latitudes in winter. ISME J. 7, 1669–1677 (2013).

16. Miravet-Verde, S., In Preparation.

17. Schmidt, T. S. B. et al. SPIRE: a Searchable, Planetary-scale mIcrobiome REsource. Nucleic Acids Res. 52, D777–D783 (2024).

18. Ruscheweyh, H.-J. et al. Cultivation-independent genomes greatly expand taxonomic-profiling capabilities of mOTUs across various environments. Microbiome 10, 212 (2022).

19. Shannon, C. E. A mathematical theory of communication. Bell Syst. Tech. J. 27, 379– 423 (1948).

20. Chao, A. Nonparametric Estimation of the Number of Classes in a Population. Scand. J. Stat. 11, 265–270 (1984).

21. Meier-Kolthoff, J. P., Göker, M., Spröer, C. & Klenk, H.-P. When should a DDH experiment be mandatory in microbial taxonomy? Arch. Microbiol. 195, 413–418 (2013).

22. Reji, L., Tolar, B. B., Chavez, F. P. & Francis, C. A. Depth-Differentiation and Seasonality of Planktonic Microbial Assemblages in the Monterey Bay Upwelling System. Front. Microbiol. 11, (2020).

23. Sánchez, P. et al. Marine picoplankton metagenomes and MAGs from eleven vertical profiles obtained by the Malaspina Expedition. Sci. Data 11, 1–12 (2024).

24. Yeh, Y.-C. & Fuhrman, J. A. Contrasting diversity patterns of prokaryotes and protists over time and depth at the San-Pedro Ocean Time series. ISME Commun. 2, 36 (2022).

25. Villarino, E. et al. Global beta diversity patterns of microbial communities in the surface and deep ocean. Glob. Ecol. Biogeogr. 31, 2323–2336 (2022).

26. Wilson, B. et al. Changes in Marine Prokaryote Composition with Season and Depth Over an Arctic Polar Year. Front. Mar. Sci. 4, (2017).

27. García, F. C., Alonso-Sáez, L., Morán, X. A. G. & López-Urrutia, Á. Seasonality in molecular and cytometric diversity of marine bacterioplankton: the re-shuffling of bacterial taxa by vertical mixing. Environ. Microbiol. 17, 4133–4142 (2015).

28. Priest, T. et al. Seasonal recurrence and modular assembly of an Arctic pelagic marine microbiome. Nat. Commun. 16, 1326 (2025).

29. Raes, E. J. et al. Seasonal patterns of microbial diversity across the world oceans. Limnol. Oceanogr. Lett. lol2.10422 (2024) doi:10.1002/lol2.10422.

30. Ma, S., Zhu, W., Wang, W., Li, X. & Sheng, Z. Microbial assemblies with distinct trophic strategies drive changes in soil microbial carbon use efficiency along vegetation primary succession in a glacier retreat area of the southeastern Tibetan Plateau. Sci. Total Environ. 867, 161587 (2023).

31. Caille, C., Duhamel, S., Latifi, A. & Rabouille, S. Adaptive Responses of Cyanobacteria to Phosphate Limitation: A Focus on Marine Diazotrophs. Environ. Microbiol. 26, e70023 (2024).

32. Thompson, L. R. et al. Patterns of ecological specialization among microbial populations in the Red Sea and diverse oligotrophic marine environments. Ecol. Evol. 3, 1780–1797 (2013).

33. Palovaara, J., et al. Stimulation of growth by proteorhodopsin phototrophy involves regulation of central metabolic pathways in marine planktonic bacteria. Proc. Natl. Acad. Sci. 111, E3650–E3658 (2014).

34. Partensky, F., Hess, W. R. & Vaulot, D. Prochlorococcus, a Marine Photosynthetic Prokaryote of Global Significance. Microbiol. Mol. Biol. Rev. 63, 106–127 (1999).

35. Johnson, Z. I. et al. Niche Partitioning Among Prochlorococcus Ecotypes Along Ocean-Scale Environmental Gradients. Science 311, 1737–1740 (2006).

36. Larkin, A. A. & Martiny, A. C. Microdiversity shapes the traits, niche space, and biogeography of microbial taxa. Environ. Microbiol. Rep. 9, 55–70 (2017).

37. Qu, L., Cai, R., Hu, Z. & Wang, H. Metagenomic assemblage genomes analyses reveal the polysaccharides hydrolyzing potential of marine group II euryarchaea. Environ. Res. 209, 112865 (2022).

38. Santoro, A. E., Richter, R. A. & Dupont, C. L. Planktonic Marine Archaea. Annu. Rev. Mar. Sci. 11, 131–158 (2019).

39. Piontek, J., Hassenrück, C., Zäncker, B. & Jürgens, K. Environmental control and metabolic strategies of organic-matter-responsive bacterioplankton in the Weddell Sea (Antarctica). Environ. Microbiol. 26, e16675 (2024).

40. Ebihara, A. et al. Structuring of particle-associated bacterial communities along the extracellular polymeric substance gradient of sinking and suspended particles in an oligotrophic, subtropical region of the western North Pacific Ocean. Front. Mar. Sci. 11, 1462522 (2024).

41. Vargas, L. C., Faria, L. C., Pereira, L. T. & Signori, C. N. Water masses drive the spatial and temporal distribution of marine Archaea in the northern Antarctic Peninsula. An. Acad. Bras. Ciênc. 96, e20240585 (2024).

42. Alonso-Sáez, L. et al. Role for urea in nitrification by polar marine Archaea. Proc. Natl. Acad. Sci. 109, 17989–17994 (2012).

43. Doytchinov, V. V. & Dimov, S. G. Microbial Community Composition of the Antarctic Ecosystems: Review of the Bacteria, Fungi, and Archaea Identified through an NGS-Based Metagenomics Approach. Life 12, 916 (2022).

44. Santoro, A. E., Kellom, M. & Laperriere, S. M. Contributions of single-cell genomics to our understanding of planktonic marine archaea. Philos. Trans. R. Soc. B Biol. Sci. 374, 20190096 (2019).

45. Danovaro, R. Understanding marine biodiversity patterns and drivers: The fall of Icarus. Mar. Ecol. e12814 (2024) doi:10.1111/maec.12814.

46. Chaudhary, C., Saeedi, H. & Costello, M. J. Bimodality of Latitudinal Gradients in Marine Species Richness. Trends Ecol. Evol. 31, 670–676 (2016).

47. Antão, L. H. et al. Temperature-related biodiversity change across temperate marine and terrestrial systems. *Nat*. Ecol. Evol. 4, 927–933 (2020).

48. Paoli, L. et al. Biosynthetic potential of the global ocean microbiome. Nature 607, 111–118 (2022).

49. Kang, D. D. et al. MetaBAT 2: an adaptive binning algorithm for robust and efficient genome reconstruction from metagenome assemblies. PeerJ 7, e7359 (2019).

50. Parks, D. H., Imelfort, M., Skennerton, C. T., Hugenholtz, P. & Tyson, G. W. CheckM: assessing the quality of microbial genomes recovered from isolates, single cells, and metagenomes. Genome Res. 25, 1043–1055 (2015).

51. Eren, A. M. et al. Anvi’o: an advanced analysis and visualization platform for ‘omics data. PeerJ 3, e1319 (2015).

52. Dmitrijeva, M. et al. The mOTUs online database provides web-accessible genomic context to taxonomic profiling of microbial communities. Nucleic Acids Res. 53, D797–D805 (2025).

53. Boyer, T. et al. World Ocean Atlas 2018. (2018).

54. Melin, F. gims - viirs Monthly climatology sea surface Chlorophyll-a concentration (4km) in mg.m^-3. European Commision, Joint Research Centre (JRC). (2018).

55. Melin, F. GIMS - SeaWiFS Monthly climatology primary production (9km) in gC.m^-2.day^-1. European Commission, Joint Research Centre (JRC). (2013).

56. SSALTO/DUACS. User Handbook: Eddy Kinetic Energy (EKE) Monthly Mean Products. AVISO+. Those products were processes by SSALTO/DUACS and distributed by AVISO+ (https://www.aviso.altimetry.fr) with support from CNES. (2021).

57. LOCEAN/CLS/CTOH/CNES. FSLE - Finite-Size Lyapunov Exponents and Orientations of the associated eigenvectors: Version 2021. CNES 10.24400/527896/A01-2022.002 (2021).

58. Stramski, D. et al. Relationships between the surface concentration of particulate organic carbon and optical properties in the eastern South Pacific and eastern Atlantic Oceans. Biogeosciences 5, 171–201 (2008).

59. Gilbert, J. A. et al. Defining seasonal marine microbial community dynamics. ISME J. 6, 298–308 (2012).

60. Treusch, A. H. et al. Seasonality and vertical structure of microbial communities in an ocean gyre. ISME J. 3, 1148–1163 (2009).

61. Rodrigues, J. F. M. et al. The MicrobeAtlas database: Global trends and insights into Earth’s microbial ecosystems. 2025.07.18.665519 Preprint at 10.1101/2025.07.18.665519 (2025).

