## Supplementary Figures for "Variations in the latitudinal diversity gradients of the ocean microbiome"

### Supplementary Material

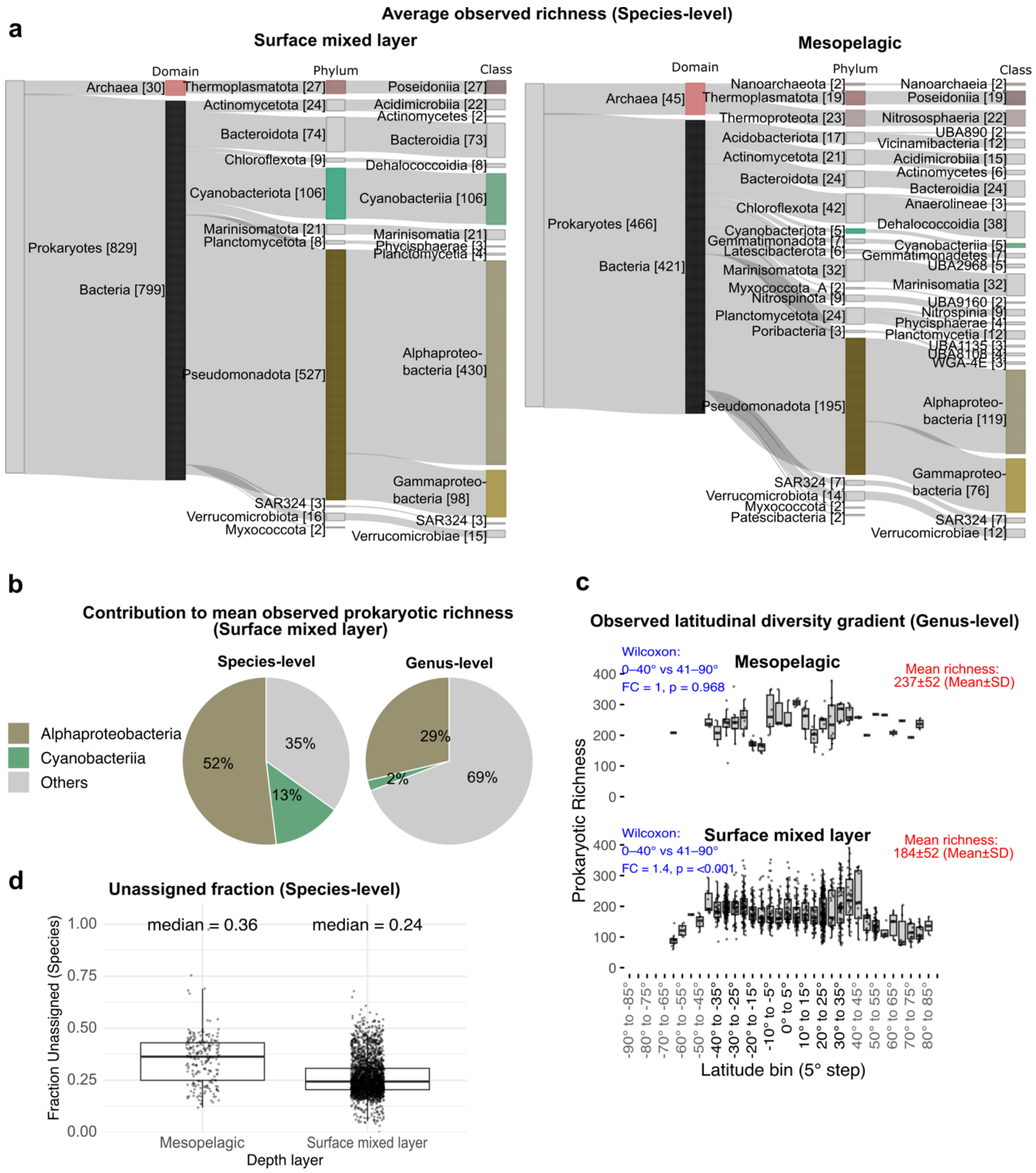


**Supplementary Fig. 1 - Distribution of prokaryotic richness across the mixed layer and mesopelagic zone.**

**(a)** Observed species-level richness of taxonomic groups, averaged across all mixed layer (left) and mesopelagic (right) samples. The Sankey graph shows all clades with mean richness > 2 for which richness distributions could be modeled or that were detected in > 50% of the mixed layer or mesopelagic samples. **(b)** Contribution of classes Alphaproteobacteria (red) and Cyanobacteriia (green) to genus-level richness (left) and species-level richness (right). Mean observed prokaryotic richness was calculated across all mixed-layer samples at both the species and genus levels. The average richness contributed by Alphaproteobacteria and Cyanobacteriia was calculated separately, and their relative contribution to total prokaryotic richness is shown in the pie charts. **(c)** Observed within-sample genus-level richness of Bacteria in the mesopelagic zone (left) and the mixed layer (right). Statistical comparisons (blue) show fold changes (FC) and p-values (Wilcoxon test) between samples from absolute latitude bins 0°–40° and 41°–90°. The top-right plots compare genus-level richness between Bacteria and Archaea. Tables report fold changes between latitude bins based on Shannon and Chao1 diversity metrics. **(d)** Sample-wise proportion of species counts that could not be assigned to an mOTU, split by mesopelagic and mixed layer.

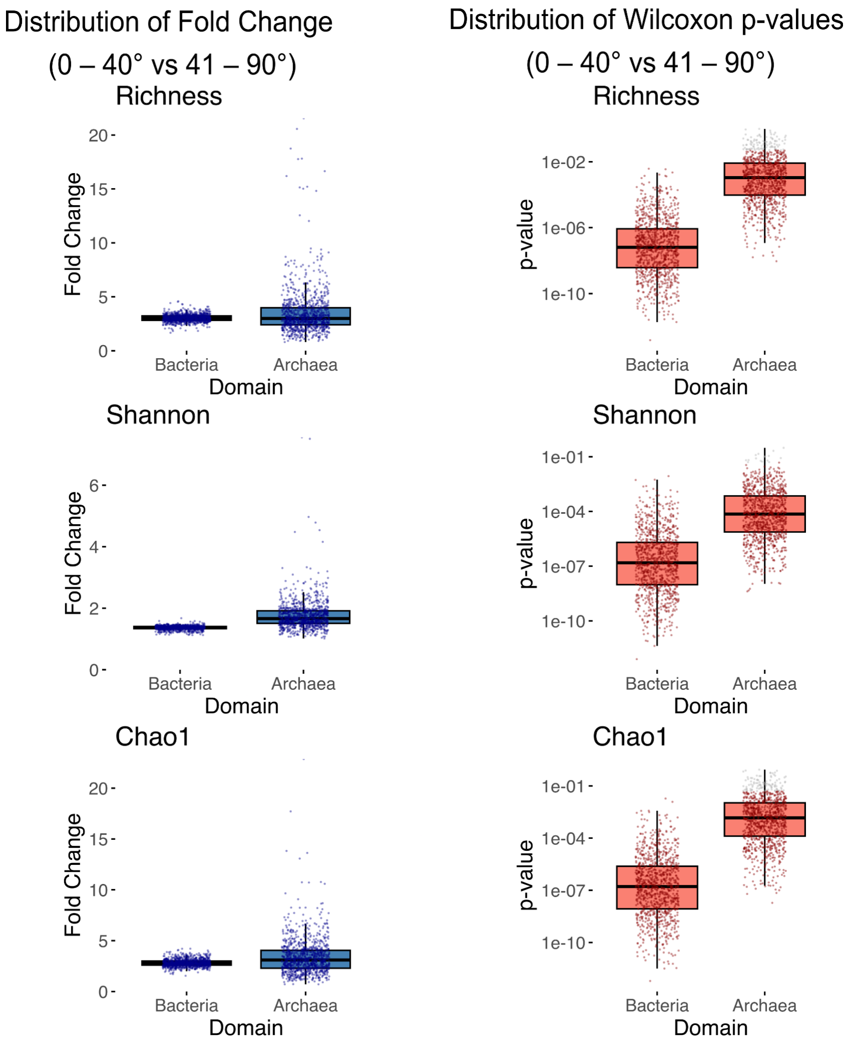


**Supplementary Fig. 2 - Robustness of mixed-layer latitudinal richness distribution.** Mixed-layer samples were randomly subsampled 1,000 times to match the number of mesopelagic samples with richness information (n = 169). For each subsample, fold changes (left) and Wilcoxon test p-values (right) were computed by comparing sample-level diversity values between the low (0°–40°) and high (41°–90°) absolute-latitude groups, separately for Bacteria and Archaea, and for richness, Shannon index, and Chao1. Points show p-values for individual subsamples and are colored red when $p < 0.05$.

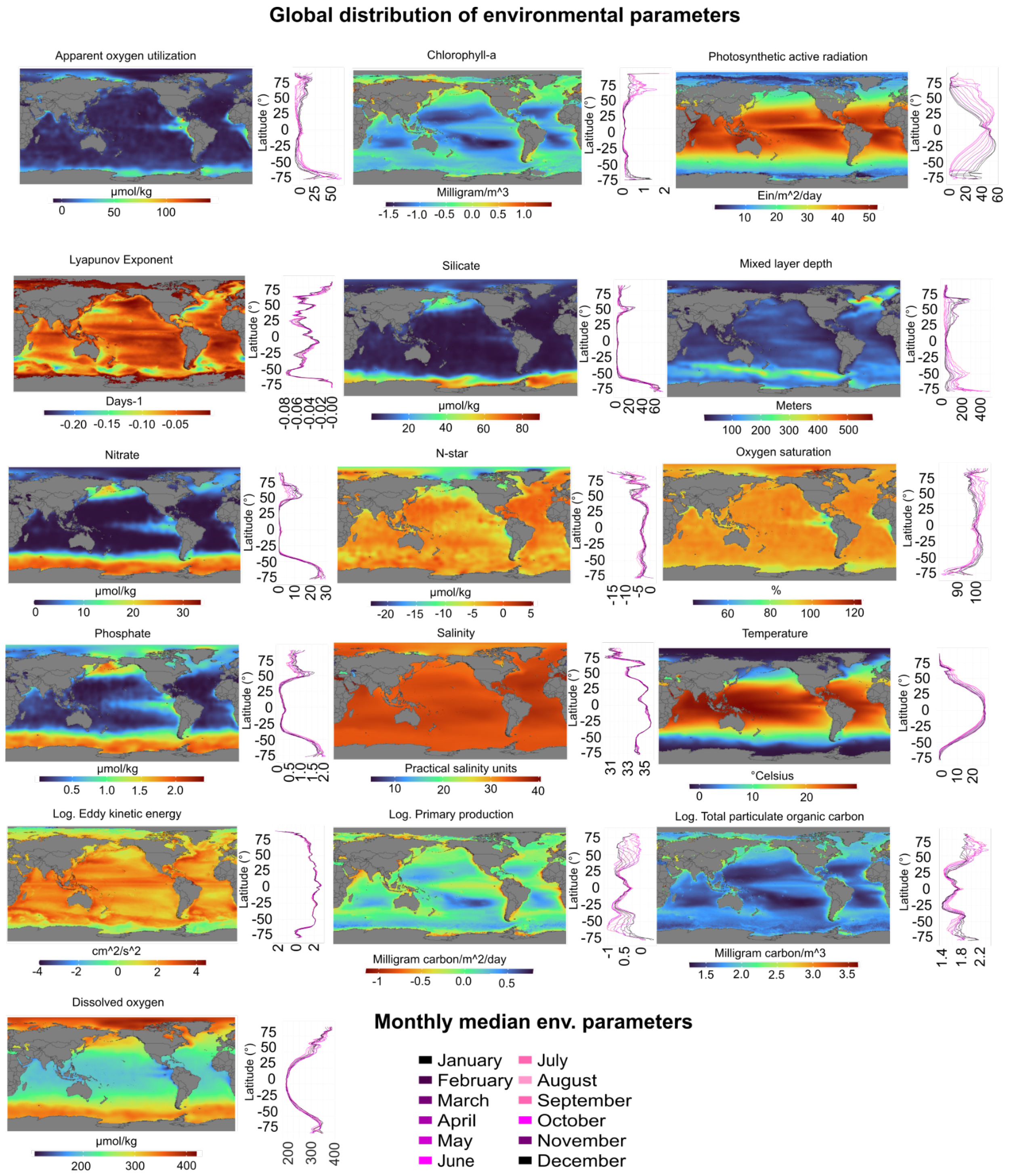


**Supplementary Fig. 3 - Global maps of environmental parameters.**
Global maps of annual mean environmental parameters in the surface ocean, centered on the Pacific basin. Each panel represents an environmental variable, including nutrient concentrations (e.g., nitrate, phosphate), chlorophyll-a, oxygen saturation, and other physical and biological parameters. Marginal line plots show seasonal variations of environmental parameters based on per 1° latitude bin, with each line representing a month.

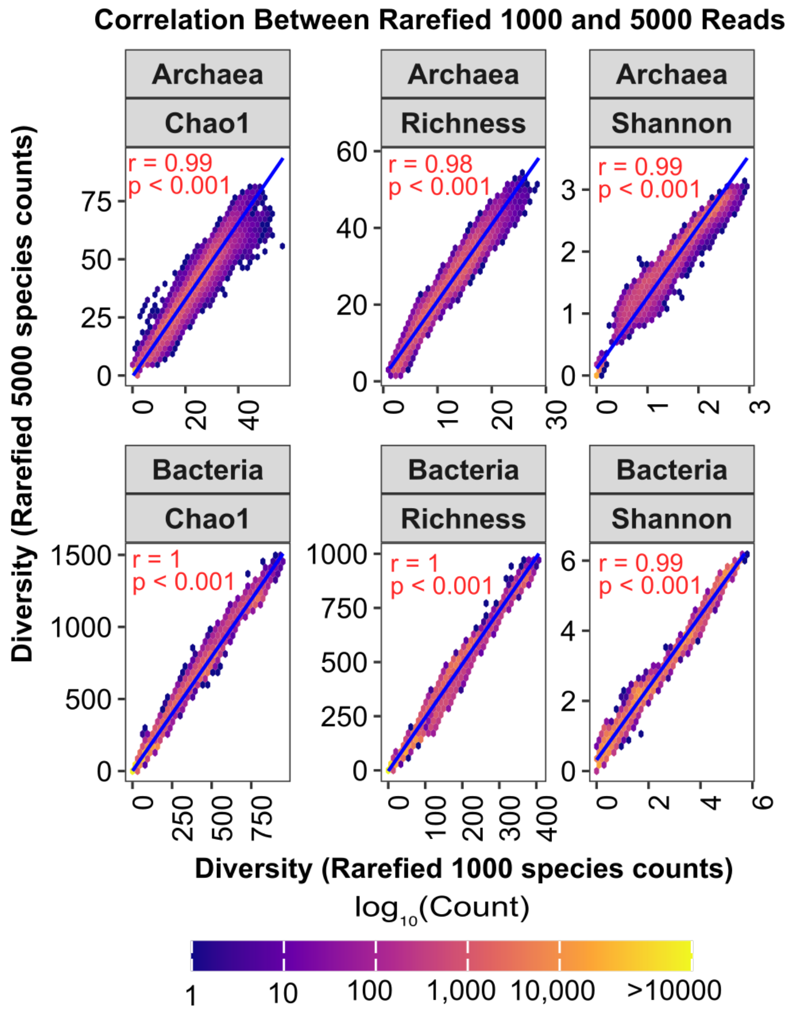


**Supplementary Fig. 4 - Comparing diversity indices across rarefaction depths in habitat models of the surface mixed layer.**

Hexbin plots comparing modeled richness, Shannon index, and Chao1 between two rarefaction levels (1,000 and 5,000 species counts) for Bacteria and Archaea. A linear fit (blue line) visualizes the relationship between the two rarefaction levels. Red text labels represent the Pearson correlation coefficients (r), and p-values quantify the statistical significance of the correlation.

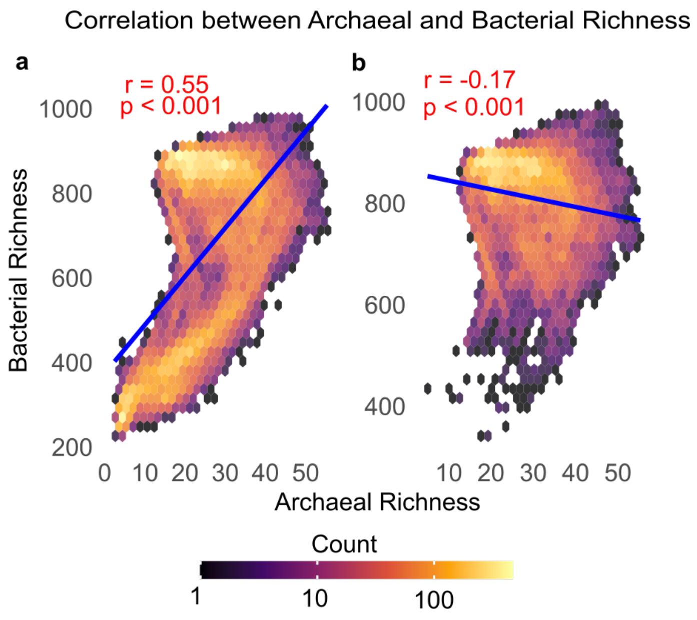


Pearson's r between bacterial and archaeal diversity

| | Shannon | Chao1 |
|---|---|---|
| Global (0-90°) | 0.86*** | 0.68*** |
| (Sub)tropics (0-40°) | -0.04*** | 0.20*** |

**Supplementary Fig. 5 - Correlation between archaeal and bacterial richness.** Hexbin plots showing the spatial correlation between modeled annual archaeal and bacterial richness in (**a**) the global ocean (0–90° absolute latitude) and (**b**) the (sub)tropics (0–40° absolute latitude). Blue lines show linear regressions; Pearson's *r* and significance were calculated using R. Correlations for Shannon index and Chao1 richness estimates show similar patterns (***$p < 0.001$).

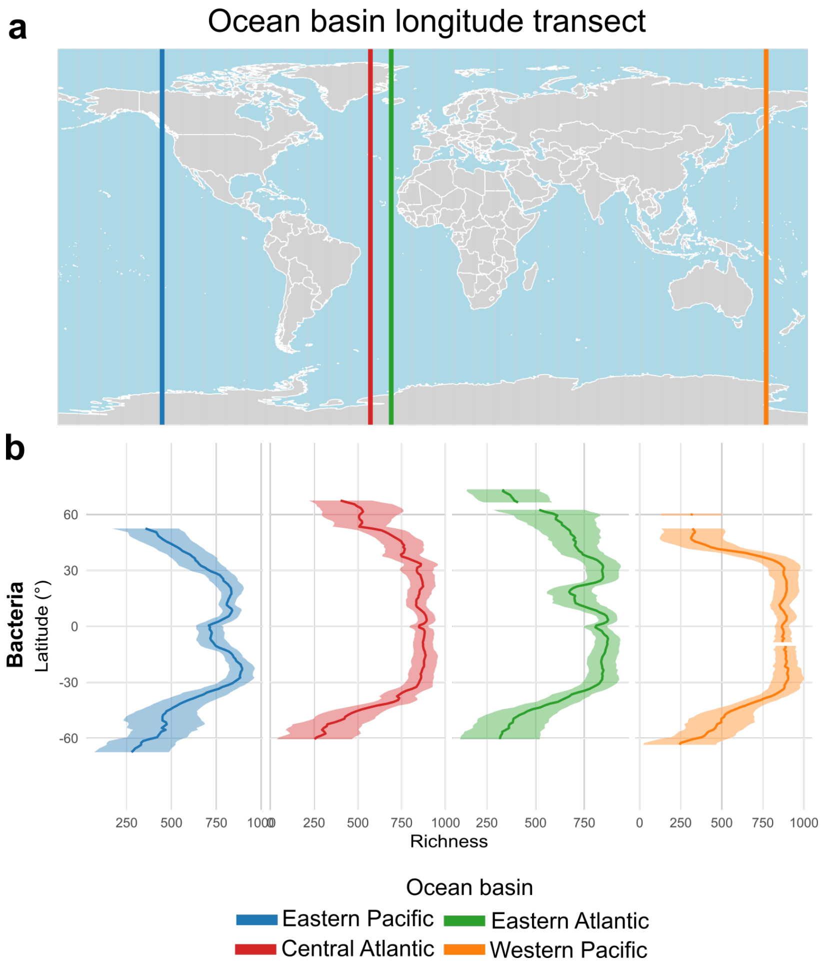


**Supplementary Fig. 6 - LDGs of marine bacteria across ocean basins.**

**(a)** Global map showing four longitudinal transects: Eastern Pacific (-130°, blue), Central Atlantic (-30°, red), Eastern Atlantic (20°, green), and Western Pacific (160°, orange). **(b)** Bacterial richness patterns along latitudinal gradients for each ocean basin.

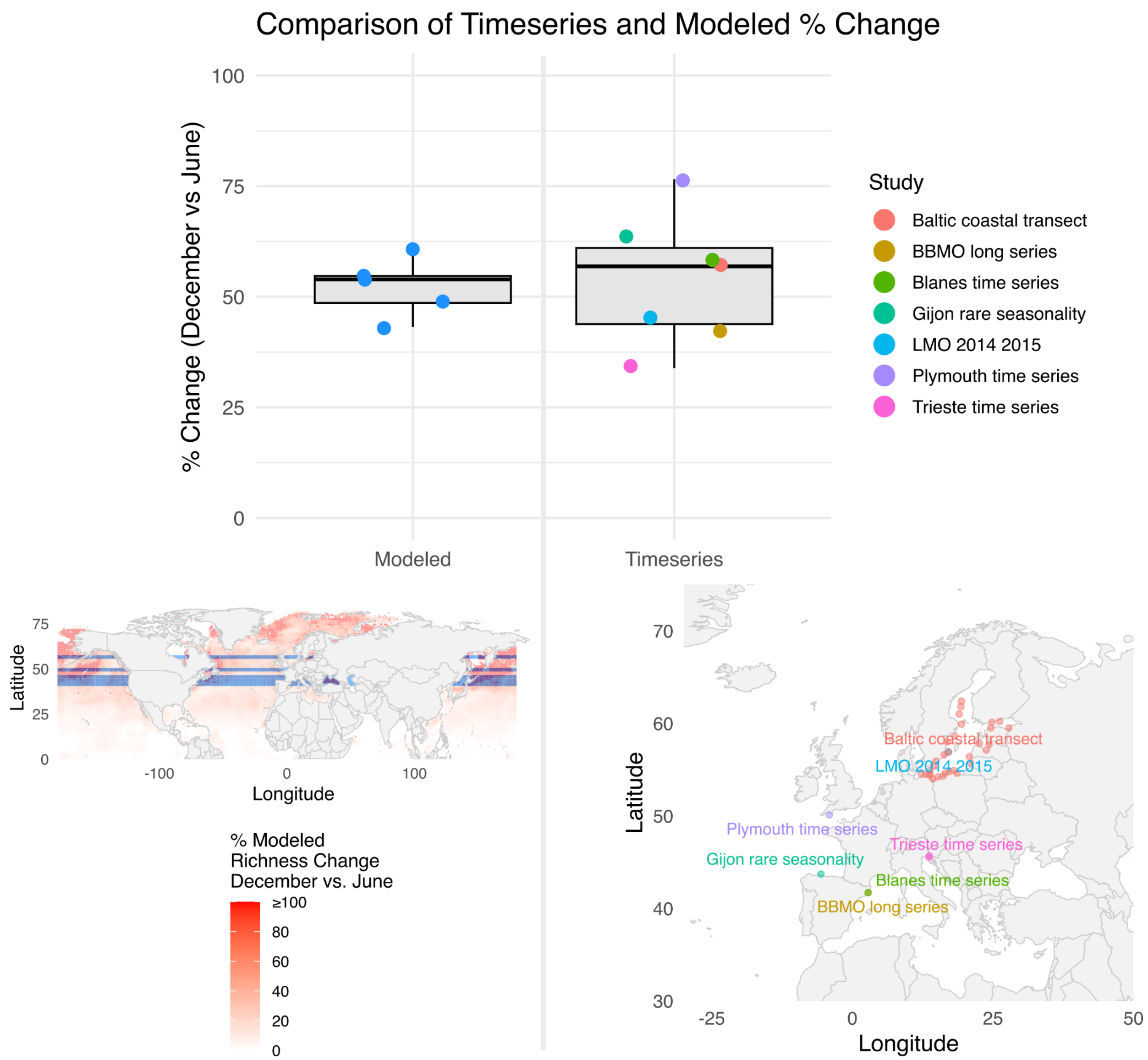


**Supplementary Fig. 7 - Boxplot of seasonal percentage increase in prokaryotic richness.** Boxplots comparing the percentage change in prokaryotic richness in December relative to June. The right panel shows observed richness from European time-series studies, and the left panel shows median modeled richness per latitude bin corresponding to the European time-series data. Maps display the sampling locations of the European time-series studies (right) and their corresponding modeled latitudes (left).

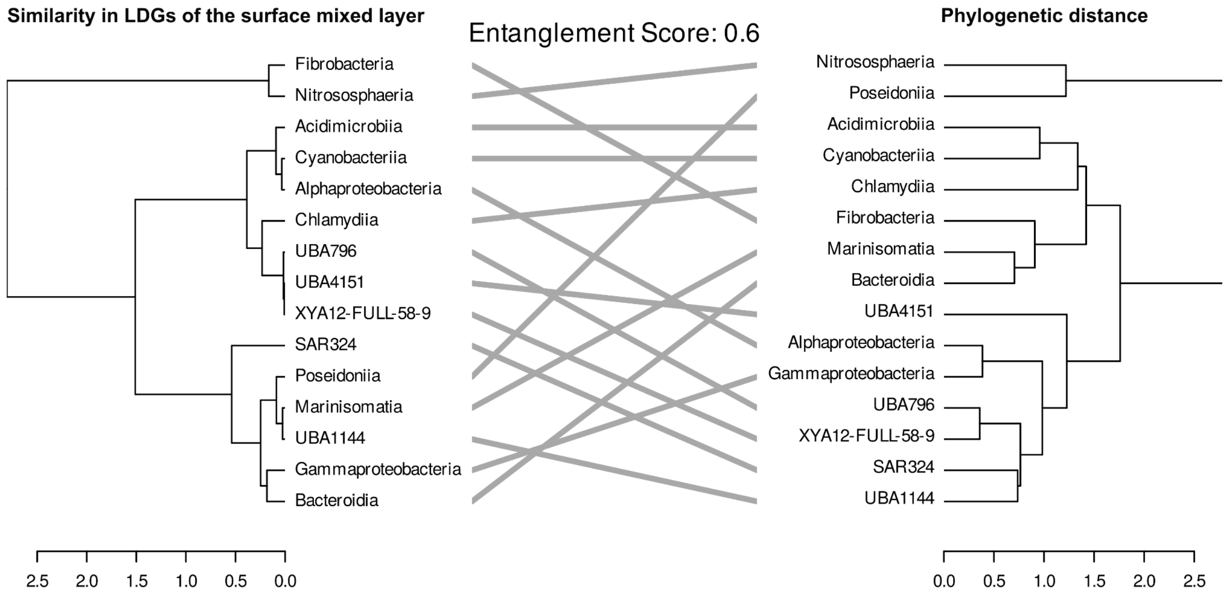


**Supplementary Fig. 8 - Comparison of similarity in LDGs and phylogenetic distances.** The left dendrogram represents the similarity of annual LDGs of the surface mixed layer (pairwise Pearson correlation, clustering algorithm “ward.D2”). The right dendrograms represent the patristic distances between modeled taxonomic ranks (GTDB version 220, clustering algorithm “ward.D2”). For each tree (Archaea and Bacteria), the respective full GTDB tree was reduced to include only nodes corresponding to modeled taxonomic groups. For calculating the entanglement score, the bacterial and archaeal trees were concatenated. The score quantifies the degree of entanglement from 0 (high alignment) to 1 (fully entangled).

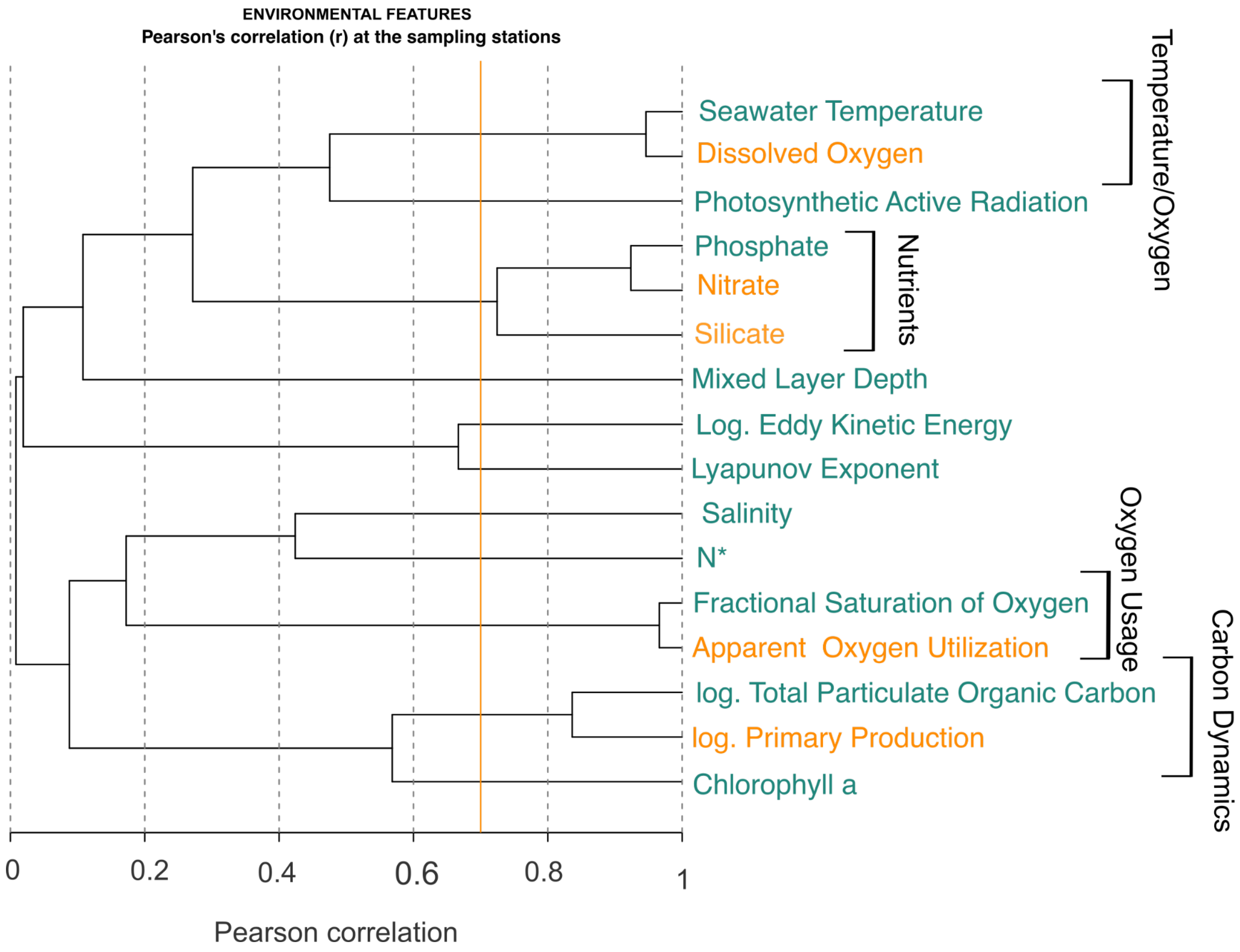


**Supplementary Fig. 9 - Multicollinearity of environmental parameters in the mixed layer ocean**.

Global multicollinearity structure of environmental features. The dendrogram depicts pairwise correlations among environmental features at a global spatial resolution of 1° longitude x 1° latitude. The red line indicates a Pearson correlation coefficient of 0.8. The orange line marks the threshold (Pearson's $r \geq 0.7$) above which two environmental features are considered closely correlated. For correlated pairs, the predictor with the higher mutual information and Pearson correlation with the target variable was retained (green), while the other was excluded from further analysis (orange).

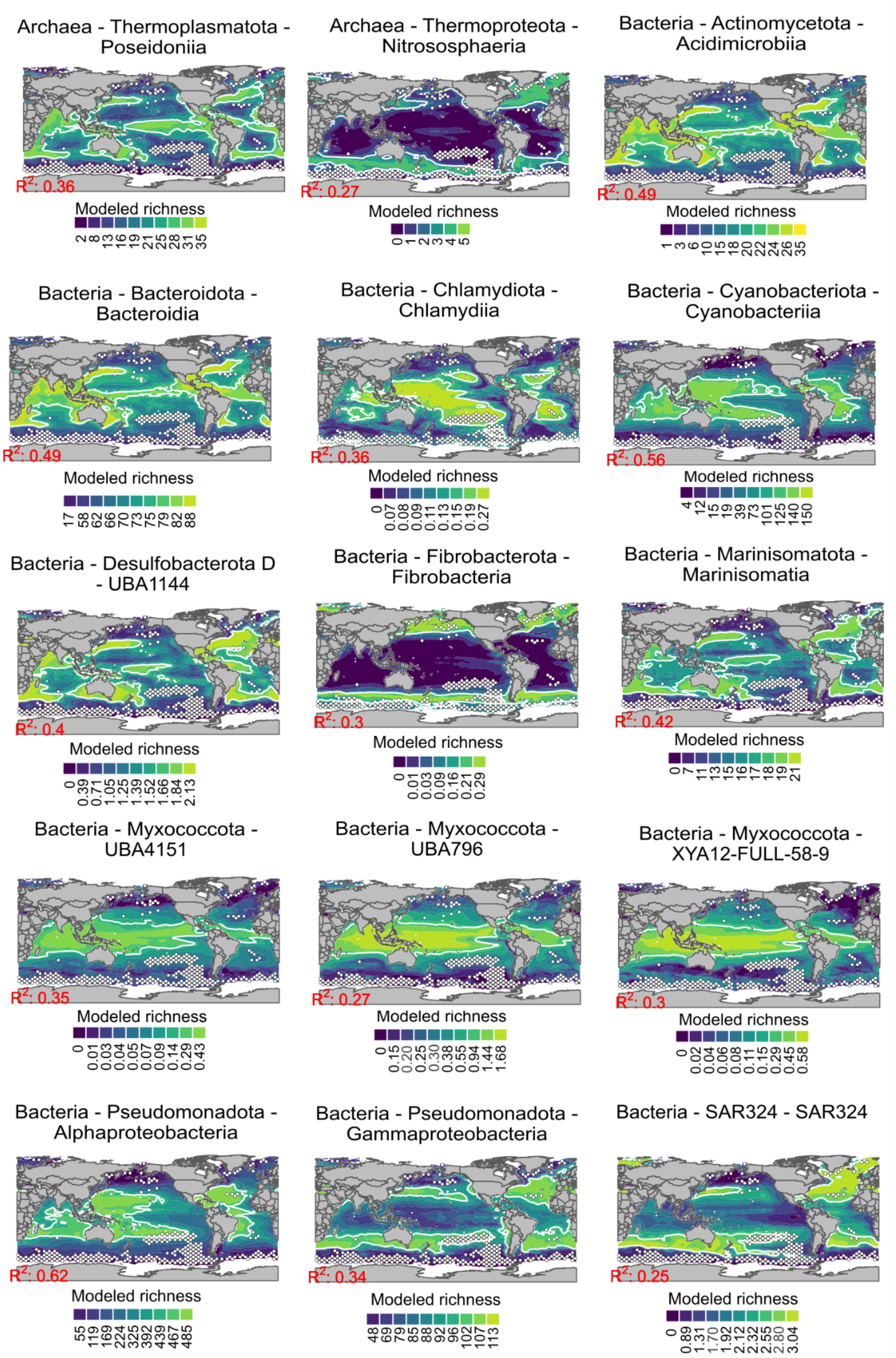


**Supplementary Fig. 10 - Global maps of annual clade-level richness.**
Global maps of class-level annual richness patterns visualized using ensemble-based predictions on a Pacific-centered world map. Areas of model uncertainty are shown as white stippling, representing regions with a coefficient of variation > 20% across ensemble members. The predictive power of each model $R^2$ is shown in red in the lower-left corner of each map. White contour lines highlight hotspots of diversity, defined as regions above the 75$^{th}$ percentile of modeled richness.

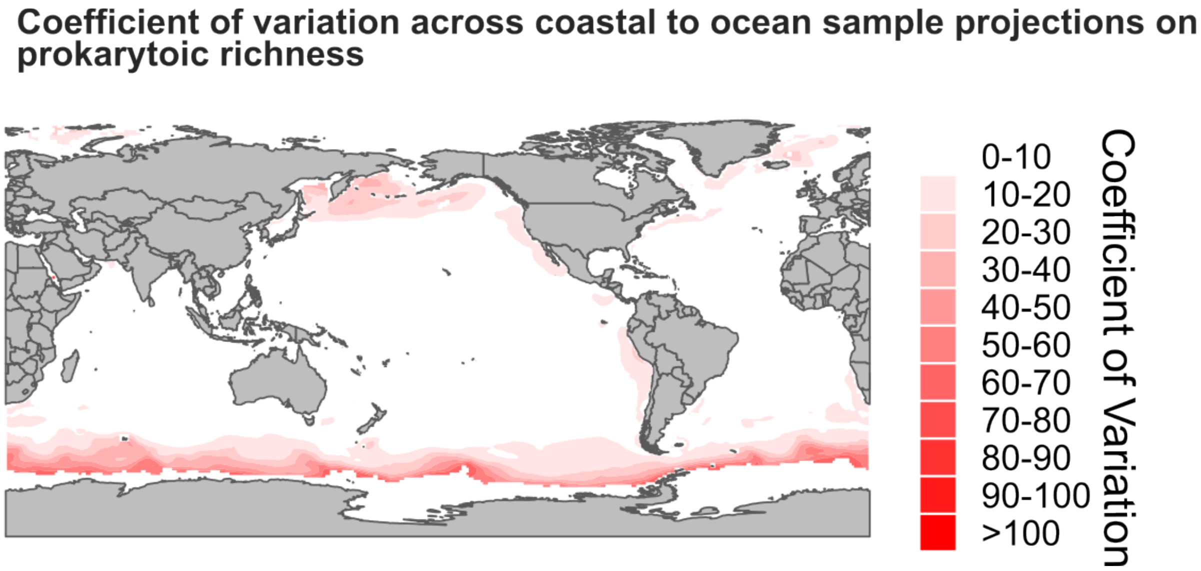


**Supplementary Fig. 11 - Effect size of coastal samples on modeled annual richness.** Coefficient of variation in predicted prokaryotic richness across global maps generated from habitat models trained on datasets with progressive exclusion of coastal samples (from 5 km to 1,500 km). The coefficient of variation is visualized using red color gradients, with darker areas indicating higher variability among models. This spatial pattern reflects the sensitivity of richness predictions to the inclusion of nearshore data.

| Study accession | Name | Latitude range (°) | Longitude range (°) | Number of samples | Amplified region |
|---|---|---|---|---|---|
| ERP117856 | Baltic_coastal_transect | 53.96–62.42 | 12.28–27.8 | 53 | (WMS) |
| ERP122219 | BBMO_long_series | 41.67 | 2.8 | 249 | V3–V4 |
| SRP091333 | Blanes_time_series | 41.67 | 2.8 | 133 | V1–V3 |
| ERP005924 | Gijon_rare_seasonality | 43.67 | -5.58 | 51 | V3–V4 |
| ERP137221 | LMO_2014_2015 | 56.93 | 17.06 | 86 | V3–V4 |
| ERP016541 | Plymouth_time_series | 50.15 | -4.13 | 146 | V6 |
| SRP364979, SRP339229, SRP364989, ERP145943 | Trieste_time_series | 45.55–45.7 | 13.55–13.71 | 186 | V4–V5 |

**Supplementary Table 1 - Metrics of European time-series studies.**
This table contains contextual information on the studies, for which taxonomic profiles were obtained from the MicrobeAtlas database.
